# Thalamocortical excitability adjustments guide human perception under uncertainty

**DOI:** 10.1101/2020.06.22.165118

**Authors:** Julian Q. Kosciessa, Ulman Lindenberger, Douglas D. Garrett

## Abstract

Adaptive human behavior builds on prior knowledge about stimulus relevance. Some environments cue such knowledge more than others. To behave adaptively, observers need to flexibly adjust sensory processing to the degree of contextual uncertainty. We hypothesize that the neural basis for these perceptual adjustments consists in the ability of the cortical network to switch back and forth between a rhythmic state that serves selective processing, and a state of elevated asynchronous neural activity that boosts sensitivity. To test this hypothesis, we recorded non-invasive EEG and fMRI BOLD dynamics while 47 healthy young adults performed a parametric visual attention task with varying numbers of relevant stimulus features. Drift-diffusion modeling of response behavior and electrophysiological signatures revealed that greater contextual uncertainty lowered the rate of evidence accumulation while increasing thalamocortical engagement, with concomitant increments in cortical excitability and pupil dilation. As predicted, uncertainty-related processing adjustments were expressed as switches between a state of phase-dependent excitability modulation in the alpha band and a state of increased irregularity of brain dynamics. We conclude that humans dynamically adjust sensory excitability according to the processing fidelity afforded by an upcoming choice, and that neuromodulatory processes involving the thalamus play a key role in adjusting excitability in the human brain.

**Highlights:**

- With increasing contextual uncertainty, human cortical networks shift from a state of phase-dependent excitability modulation in the alpha band into a state of elevated excitatory tone and asynchronous neural activity
- Evidence based on joint modeling of behavior, EEG, and BOLD suggests that neuromodulatory processes involving the thalamus regulate these shifts
- Theoretical and empirical considerations suggest contributions of both frequency-specific and aperiodic neural dynamics to human behavior

## Introduction

Adaptive behavior requires dynamic adjustments to the perception of high-dimensional inputs. Prior knowledge about the momentary relevance of specific environmental features selectively enhances their processing while suppressing distractors (for reviews see Buschman & Kastner, 2015; Desimone & Duncan, 1995; Maunsell, 2015), which can be implemented via gain modulation in sensory cortex (Ferguson & Cardin, 2020). Crucially, *a priori* information regarding feature relevance is not always available; and how the brain flexibly adjusts the processing of complex inputs according to contextual uncertainty remains unclear (Bach & Dolan, 2012).

Selective gain control has been associated with ***phasic*** (i.e., phase-dependent) inhibition of task-irrelevant stimulus dimensions during cortical alpha (^~^8-12 Hz) rhythms (Klimesch, Sauseng, & Hanslmayr, 2007; Sadaghiani & Kleinschmidt, 2016). In particular, rhythmic modulations of feedforward excitability (Haegens, Nacher, Luna, Romo, & Jensen, 2011; Lorincz, Kekesi, Juhasz, Crunelli, & Hughes, 2009) may provide temporal ‘windows of opportunity’ for high-frequency gamma synchronization in sensory cortex (Spaak, Bonnefond, Maier, Leopold, & Jensen, 2012; van Kerkoerle et al., 2014) and increased sensory gain (Fries, 2015; Ni et al., 2016; Peterson & Voytek, 2017). However, specifically increasing the fidelity of single stimulus dimensions is theoretically insufficient when uncertain environments require joint sensitivity to multiple stimulus features (Pettine, Louie, Murray, & Wang, 2020). During high uncertainty, transient increases to the ***tonic*** excitation/inhibition (E/I) ratio in sensory cortex provide a principled mechanism for elevated sensitivity to – and a more faithful processing of – high-dimensional stimuli (Destexhe, Rudolph, & Pare, 2003; Marguet & Harris, 2011). In electrophysiological recordings, scale-free 1/f slopes are sensitive to differences in E/I ratio (Gao, Peterson, & Voytek, 2017), and vary alongside sensory stimulation (Billig et al., 2019; Podvalny et al., 2015) and arousal states (Colombo et al., 2019; Lendner et al., 2019). Whether contextual demands modulate scale-free activity is unknown however. We hypothesize that high uncertainty shifts cortical regimes from rhythmic excitability modulations towards tonic excitability increases.

Such state switches in network excitability may be shaped by neuromodulation and subcortical activity (Harris & Thiele, 2011). Neuromodulation potently alters cortical states (Froemke, 2015; Thiele & Bellgrove, 2018) and sensory processing (Berridge & Waterhouse, 2003; McCormick, Pape, & Williamson, 1991; McGinley, David, & McCormick, 2015), and noradrenergic arousal in particular may permit high sensitivity to incoming stimuli (Posner & Rothbart, 2007). Yet, non-invasive evidence is lacking for whether/how neuromodulation affects contextual adaptability. Moreover, despite early proposals for thalamic involvement in attentional control (Crick, 2003; Jasper, 1948; Rafal & Posner, 1987), studies have dominantly focused on cortical information flow (e.g., Siegel, Buschman, & Miller, 2015), at least in part due to technical difficulties in characterizing thalamic contributions. Crucially, the thalamus provides a nexus for the contextual modulation of cortical circuits (Halassa & Kastner, 2017; Honjoh et al., 2018), is a key component of neuromodulatory networks (McCormick et al., 1991; Schiff, 2008; Song et al., 2017) and robustly modulates system excitability via rhythmic and aperiodic membrane fluctuations (Jones, 2009). However, human evidence for a central thalamic role in cortical state adjustments at the service of behavioral flexibility is missing.

Here, we aimed at overcoming this lacuna by assessing the effects of contextual uncertainty during stimulus encoding on cortical excitability, neuromodulation, and thalamic activity in humans. We performed a multi-modal (parallel) EEG-fMRI experiment to capture both fast cortical dynamics (EEG) and subcortical activity (fMRI) while recording pupil dilation as a non-invasive proxy for neuromodulatory drive (Joshi & Gold, 2020). Participants performed a parametric adaptation of the classic dot motion task (Gold & Shadlen, 2007) (Figure 1). Specifically, we manipulated the number of stimulus dimensions that are task-relevant in a given trial while holding the sensory features of the task (i.e., its appearance on the screen) constant across trials. By applying drift-diffusion modeling to participants’ choice behavior while jointly assessing electrophysiological signatures of decision processes, we found that uncertainty during sensation reduces the rate of subsequent evidence integration. This reduction in available sensory evidence for single targets was associated with increased cortical excitability, as indexed by joint low-frequency (~alpha) desynchronization and high-frequency (~gamma) synchronization, and an increase in E/I ratio, as indicated by increased sample entropy and flatter scale-free 1/f slopes, during stimulus processing, in lines with broad sensitivity increases during periods of higher uncertainty. These excitability adjustments occurred in parallel with increases in pupil-based arousal. Finally, inter-individual differences in the modulation of cortical excitability, drift rates and arousal were jointly associated with the extent of thalamic BOLD signal modulation, pointing to the importance of subcortical mechanisms for cortical state adjustments. Together, these findings suggest that neuromodulatory processes involving the thalamus shape cortical excitability states in humans, and that a shift from alpha-rhythmic to aperiodic neural dynamics adjusts the processing fidelity of external stimuli in service of upcoming decisions.

**Figure 1.**
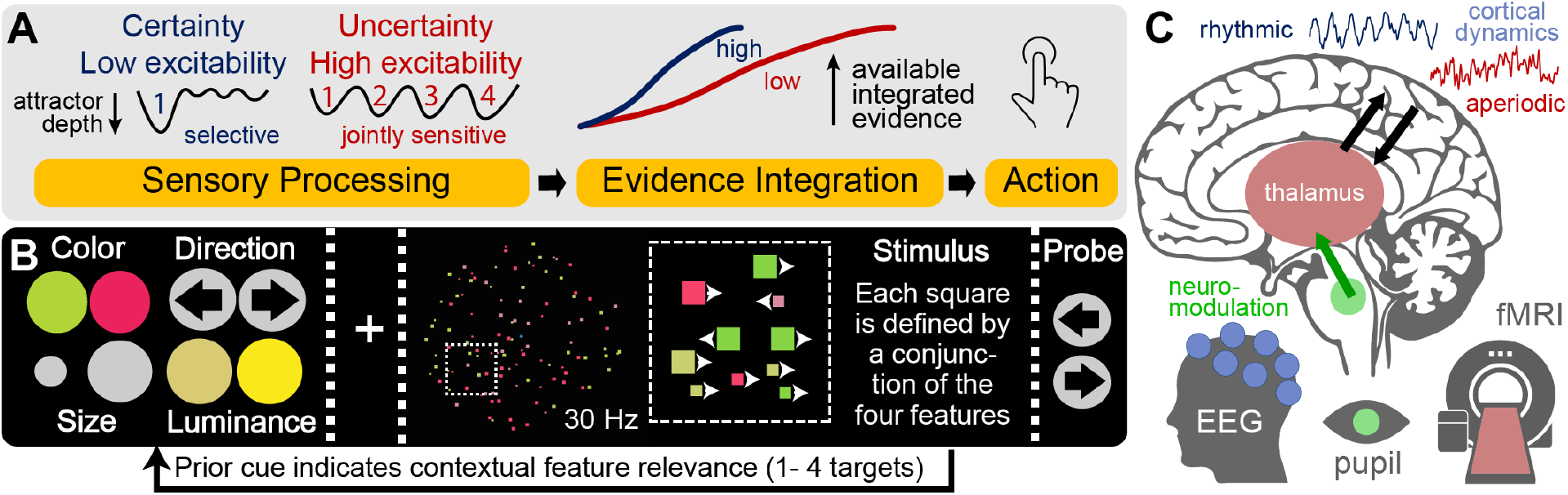
Hypotheses & task design. (**A**) We probed whether participants modulate cortical excitability during stimulus processing to guide subsequent evidence accumulation. We hypothesized that when valid attentional cues about a single target feature are available in advance, a low excitability regime may optimize subsequent choices via the targeted selection of relevant – and inhibition of irrelevant – information. This can be conceptualized as the creation of a “single feature attractor.” In contrast, under high probe uncertainty, higher excitability may afford the concurrent sampling of multiple relevant features, but at the cost of a relative reduction of subsequently available evidence for any individual feature. (**B**) Participants performed a **M**ulti-**A**ttribute **A**ttention **T**ask (“MAAT”) during which they had to sample up to four visual features in a joint display for immediate subsequent recall. Prior to stimulus presentation, participants were validly cued to a set of potential target probes. The number and identity of cues were varied to experimentally manipulate the level of expected probe uncertainty. (**C**) We hypothesized that increasing probe uncertainty would induce a joint increase in neuromodulation and thalamic activity, associated with shifts from a phasic gain control mode (implemented via neural alpha rhythms) toward transient increases in tonic excitability (as indicated by aperiodic cortical activity). Participants performed the same task in both an EEG and an fMRI session, allowing us to assess joint inter-individual differences in fast cortical dynamics (EEG) and subcortical sources (fMRI).

## Results

We developed a dynamic visual **M**ulti-**A**ttribute **A**ttention **T**ask (“MAAT”) to uncover rapid adjustments to stimulus processing and perceptual decisions under expected uncertainty (Figure 1). Participants visually sampled a moving display of small squares, which were characterized by four stimulus features, with two exemplars each: their color (red/green), their movement direction (left/right), their size (large/small), and their color saturation (high/low). Any individual square was characterized by a conjunction of the four features, while one exemplar of each feature (e.g., green color) was most prevalent in the entire display. Following stimulus presentation, participants were probed on a single feature as to which of the two exemplars was most prevalent (via 2-AFC). Probe uncertainty was parametrically manipulated using valid pre-stimulus cues, indicating the feature set from which a probe would be selected. The feature set remained constant for a sequence of eight trials to reduce set switching demands. Optimal performance required flexible sampling of the cued feature set while jointly inhibiting uncued features; participants had to thus rapidly encode a varying number of targets (“target load”) to prepare for an upcoming probe. Participants performed the task well above chance level for different features and for different levels of probe uncertainty (Figure S1A). As the number of relevant targets increased, participants systematically became slower (median RT; EEG: β = .138, *p* ~ 0; MRI: β = .107, *p* ~ 0) and less accurate (EEG: β = −.032, *p* ~ 0; MRI: β = −.025, *p* = 2.4e-07) in their response to single-feature probes (Figure S1B).

### Probe uncertainty during sensation decreases the rate of subsequent evidence integration

We leveraged the potential of sequential sampling models to disentangle separable decision processes in order to assess their modulation by probe uncertainty. In particular, drift-diffusion models estimate (a) the non-decision time (NDT), (b) the drift rate at which information becomes available, and (c) the internal evidence threshold or boundary separation (see Figure 2A; for a review see Forstmann, Ratcliff, & Wagenmakers, 2016). We fitted a hierarchical drift-diffusion model (HDDM) separately for each testing session, and assessed individual parameter convergence with established EEG signatures (Donner, Siegel, Fries, & Engel, 2009; O’Connell, Dockree, & Kelly, 2012; Twomey, Kelly, & O’Connell, 2016; van Vugt, Beulen, & Taatgen, 2019). In particular, we investigated the Centroparietal Positive Potential (CPP) and lateralized beta suppression as established neural signatures of evidence integration from eidetic memory traces (Twomey et al., 2016). The best behavioral fit was obtained by a model incorporating probe uncertainty-based variations in drift rate, non-decision time and boundary separation (Figure S1B). Yet, there was no evidence for modulation of the threshold of the CPP or the contralateral beta response (Figure S1C). In line with prior work (McGovern, Hayes, Kelly, & O’Connell, 2018), we therefore selected an EEG-informed model with fixed thresholds across target load levels. With this model, reliability of individual parameters as well as of their load-related changes was high across EEG and MRI sessions (see below and Figure S1E, F). Parameter interrelations are reported in Text S1.

**Figure 2:**
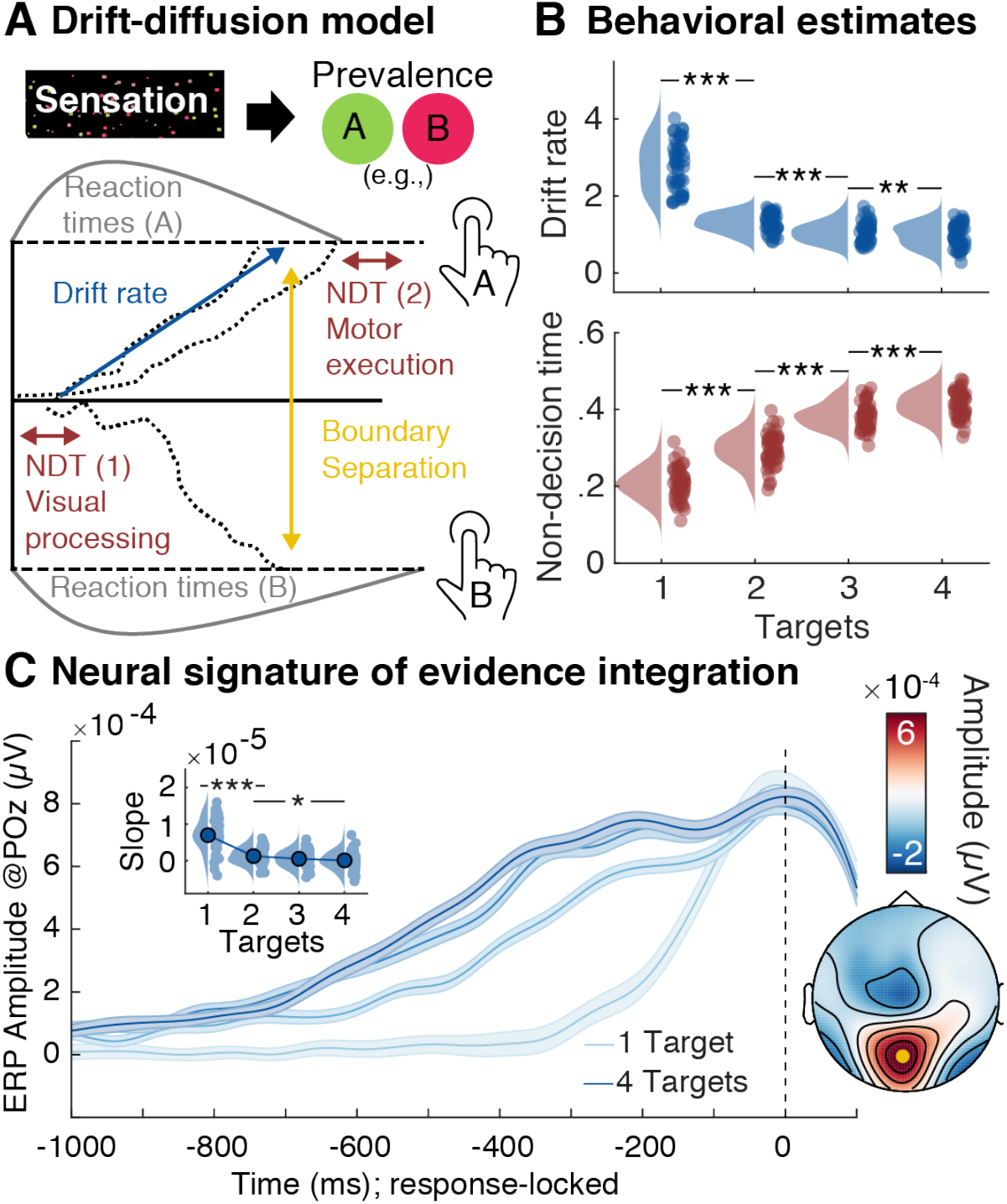
Evidence integration upon probe presentation decreases as a function of prior uncertainty. (**A**) Schematic of drift-diffusion model. Following visual encoding, evidence is successively accumulated towards either of two bounds when probed for the dominant prevalence of one of two options of a single feature. A button press indicates the decision once one of the bounds has been reached and motor preparation has concluded. A non-decision time parameter captures visual encoding and motor preparation, drift rate captures the amount of available information, and boundary separation captures response bias i.e., conservative vs. liberal). (**B**) Behavioral parameter estimates for drift rate and non-decision time (NDT; discussed in Text S3), as indicated by the hierarchical drift-diffusion model (HDDM). (**C**) Modulation of the Centroparietal Positive Potential (CPP) as a neural signature of evidence accumulation (mean +− within-subject SE). The probe-locked CPP indicates decreases in drift rate with prior probe uncertainty. Insets show CPP slope estimates from −250 to −100 ms relative to response execution, as well as the corresponding topography (CPP channel shown in yellow). [*** p < .001, ** p < .01, * p < .05]

Behavioral model estimates (Figure 2B) and EEG signatures (Figure 2C, Figure S2A) jointly indicated that probe uncertainty during stimulus presentation decreased the drift rate during subsequent evidence accumulation. This indicates a reduction of available evidence for single features when more features had to be sampled. Individual drift rate estimates for a single target were positively correlated with the slope of the CPP (r = 0.52, 95%CI [0.26, 0.71], p = 3.59e-4), while individual drift rate reductions reflected the shallowing of CPP slopes (r(137) = 0.34, 95%CI [0.18, 0.48], p = 4.87e-5). Notably, the magnitude of evidence decreases with increasing probe uncertainty was strongly anticorrelated with the available evidence when the target attribute was known in advance (i.e., the single target condition; EEG session: *r* = −.93, *p* = 4e-22, MRI session: *r* = −.88, *p* = 1e-15). That is, participants with more available evidence after selectively attending to a single target showed larger drift rate decreases under increased probe uncertainty. Importantly however, participants with higher drift rates for single targets also retained higher drift rates at higher probe uncertainty (i.e., high reliability for e.g., four targets: EEG: r = .48; p = 6e-4; MRI: r = .53, p = 2e-4). Moreover, individuals with higher drift rates across target loads exhibited lower average RTs (EEG: *r* = −.42, *p* = .003; MRI: *r* = −.41, *p* = .007) and higher task accuracy (EEG: *r* = .86, *p* = 2e-14; MRI: *r* = .89, *p* = 4e-16). Thus, in the present paradigm, more pronounced drift rate decreases with increasing probe uncertainty index a successful modulation of feature-based attention during encoding, and better overall performance.

We performed multiple control analyses to further elucidate decision properties. First, we did not observe a similar ramping of the CPP during stimulus presentation (Figure S2B), suggesting that evidence accumulation was primarily initiated by the probe. Second, drift rate reductions were not primarily driven by differences between feature attributes (Figure S2C). Third, concurrent variations in response agreement across cued attributes could not account for the observed effects (Text S2; Figure S1D). Fourth, individual drift rates for single targets were unrelated to threshold estimates (EEG: *r* = −.005, *p* = .74; MRI: *r* = −.006; *p* = .72), thus suggesting a lack of differences in response bias (Ratcliff & McKoon, 2008). Finally, participants with larger drift rate decreases exhibited more constrained non-decision time increases (EEG: r(137) = 0.32, 95% CI [0.16, 0.47], p = 1.04e-4; MRI: r(122) = 0.37, 95%CI [0.2, 0.51], p = 2.48e-5), indicating reduced additional motor transformation demands (see Text S3) in high performers.

### Cortical excitability increases under uncertainty guide subsequent evidence integration

Decreases in the rate of evidence integration indicate the detrimental consequences of probe uncertainty, but not the mechanisms by which sensory processing is altered. To investigate the latter, we examined rhythmic and aperiodic cortical signatures during stimulus processing. To jointly assess multivariate changes in spectral power as a function of probe uncertainty, we performed a partial-least-squares (PLS) analysis that produces low-dimensional, multivariate relations between brain-based data – in this case time-frequency-space matrices – and other variables of interest (see methods). First, we assessed evoked changes compared to baseline using a task PLS. We observed a single latent variable (LV; permuted *p* < .001) with jointly increased power in the delta-theta and gamma bands and decreased alpha power upon stimulus onset (Figure S3A, Figure S4A), in line with increased cognitive control (Cavanagh & Frank, 2014) and heightened bottom-up visual processing (van Kerkoerle et al., 2014). We next performed a task PLS to assess spectral power changes as a function of target load. A single LV (permuted *p* < .001; Figure 3) indicated a stronger expression of this control- and excitability-like pattern with increasing probe uncertainty. Next, we assessed the link between individual changes in multivariate loadings on this “spectral power modulation factor” (SPMF) and behavioral modulations. We performed *partial repeated measures correlations (*see methods), a mixed modelling approach that controls for the main effect of probe uncertainty in both variables of interest and indicates interindividual associations independent of the specific shape of condition modulation in individual participants. Crucially, individual SPMF loadings were positively correlated with interindividual performance differences during selective attention (Figure 3F) and uncertainty-related performance changes (Figure 3G). Participants with stronger spectral power modulation during sensation exhibited faster evidence integration in the selective attention condition, as well as a stronger drift rate decreases under uncertainty [r(137) = −0.4, 95%CI [−0.53, −0.25], p = 1.12e-6], while showing constrained increases in non-decision time [r(137) = −0.26, 95%CI [−0.41, −0.1], p ~ 0]. In sum, this suggests that high performers flexibly increased visual throughput as more features became relevant via top-down control of cortical excitability.

**Figure 3:**
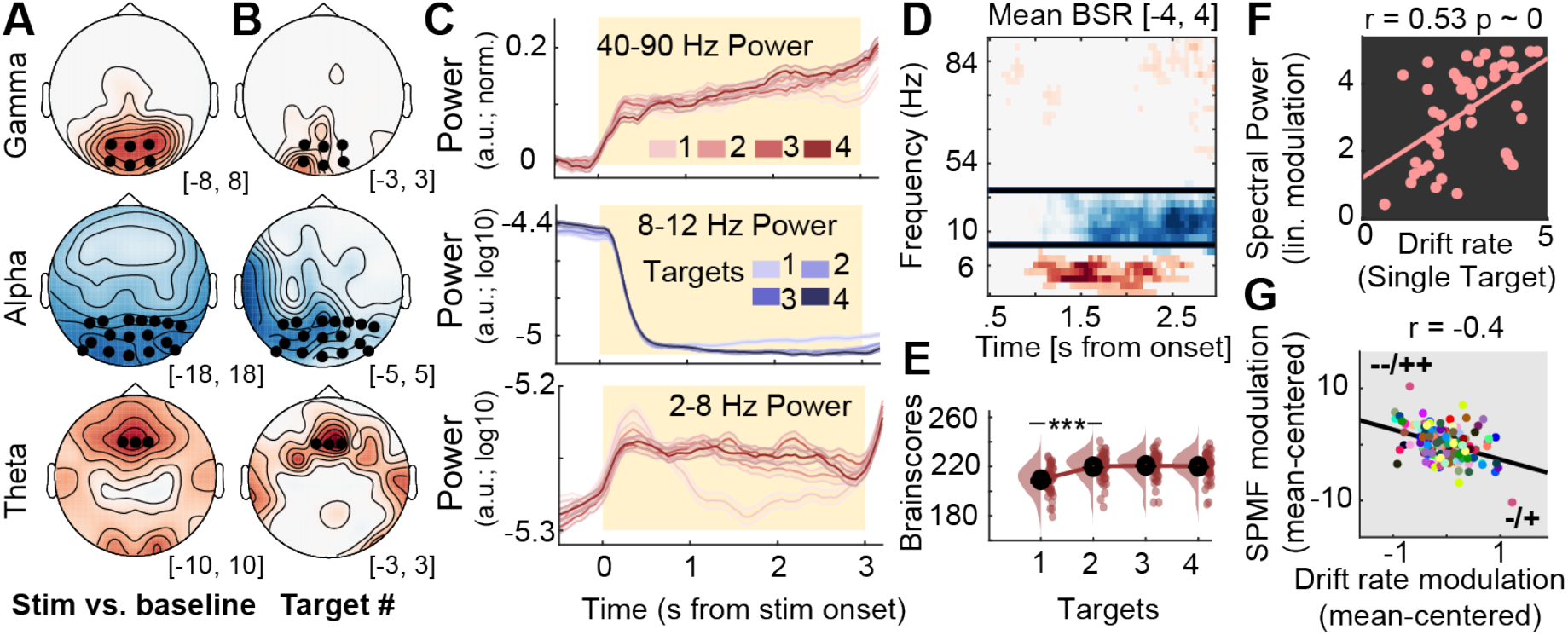
Multivariate power changes with probe uncertainty during stimulus encoding. (**A, B**) Topographies of stimulus-evoked power changes relative to pre-stimulus baseline (A, see Figure S3-1) and load-related power modulation (B). With increasing attentional demands, theta and ‘broadband’ gamma power increased, whereas alpha rhythms desynchronize. Asterisks indicate the sensors across which data were averaged for presentation in D. Values indicate maximum (theta/gamma) or minimum (alpha range) bootstrap ratios (BSR) across time in the clusters. (**C**) Temporal traces of band-limited power as a function of target load, extracted from the clusters presented in D (mean +− within-subject SE). (**D, E**) Multivariate loading pattern (D) for spectral power changes under uncertainty and associated multivariate brain scores at different levels of target load (E). Black bars in panel D indicate discrete frequency ranges or sensors (shown in A). (**F, G**) Participants with stronger multivariate power modulation exhibit stronger drift rates for single targets (F), as well as stronger drift rate decreases under uncertainty (G). In G, dots represent linear model residuals (see methods), colored by participant. Coupled changes across target conditions are indicated by the black line. We indicate the direction of main effects for each variable via + and − (− = small decreases, −− = large decreases, + = small increases, ++ = large increases), with directions of variables on the x-axis indicated first. [*** p < .001]

Here too, we performed multiple control analyses. First, the same multivariate power-band relations noted in our task PLS model (SPMF above) were also identified in a behavioral PLS model intended to estimate optimal statistical relations between power bands and behavior (Text S4, Figure S4B). Second, while we observed increases in pre-stimulus alpha power with increasing probe uncertainty, these changes did not relate to behavioral changes or power changes during stimulus processing (Text S5, Figure S4C). Third, the entrained steady-state visual evoked potential (SSVEP) magnitude was not modulated by target load (Text S6, Figure S4D). Fourth, multivariate power changes corresponded to narrow-band, rhythm-specific indices in the theta and alpha band (Text S7, Figure S4E), and thus did not exclusively result from changes in the aperiodic background spectrum (see below).

### Alpha phase modulates gamma power during sensation

Alpha rhythms have been related to phasic control over bottom-up input, as putatively encoded in gamma power (Spaak et al., 2012). To assess phase-amplitude coupling (PAC) in the present data, we selected temporal alpha episodes at the single-trial level (see methods, Figure 4A) and assessed the coupling between alpha phase and gamma power. We observed significant alpha-gamma PAC (Figure 4B, D left), consistent with alpha-phase-dependent excitability modulation. This was constrained to the occurrence of alpha episodes, as no significant alpha-gamma PAC was observed prior to indicated alpha episodes (grey shading in Figure 4A; Figure 4D right). Phasic gamma power modulation was observed across target load levels (Figure 4F), but alpha duration decreased as a function of load (Figure 4C). This suggests that alpha rhythms consistently regulated gamma power, but that alpha engagement decreased as more targets became relevant.

**Figure 4.**
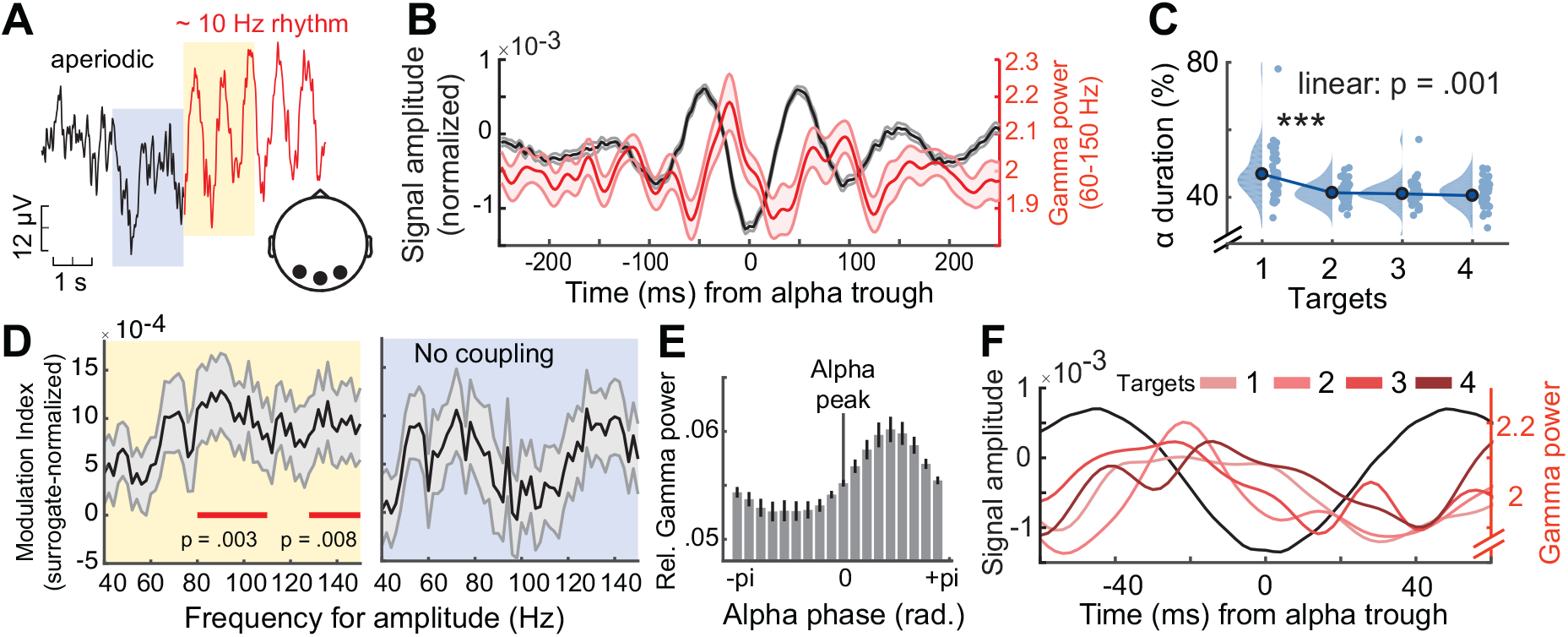
Alpha phase modulates gamma power during sensation. (**A**) Exemplary time series around the onset of a detected alpha event (example from 4-target condition). Segments were pooled across occipital channels (black dots in inset topography) and target load conditions. (**B**) Normalized gamma power (red; mean +− SE) during alpha events (yellow shading in A), is modulated by alpha phase (see methods). The unfiltered ERP aligned to the alpha trough is shown in black. Shaded regions indicate standard errors. (**C**) The relative duration of alpha events decreased with increased feature relevance. Data are individually centered across target loads. (**D**) Modulation index (MI) indicated significant coupling between the phase of alpha and gamma power during rhythmic events (left), but not during periods immediately prior to rhythm onset (right). MI was normalized using surrogate data to reduce erroneous coupling (see methods). Shaded regions indicate standard errors. (**E**) Gamma power (averaged from 60-150 Hz; mean +− SE) was maximal following alpha peaks. Power was normalized across all phase bins (see methods). (**F**) Gamma power systematic peaks between the peak and trough of alpha rhythms across target levels. For this analysis, alpha events were collapsed across all participants. [*** p < .001]

### Sample entropy and scale-free dynamics indicate shifts towards increased excitability

Next, we assessed whether reduced alpha engagement was accompanied by increases in temporal irregularity, a candidate signature for system excitability (Kosciessa, Kloosterman, & Garrett, 2020). We probed time-resolved fluctuations in sample entropy (SampEn), an information-theoretic estimate of signal irregularity. As sample entropy is jointly sensitive to broadband dynamics and narrowband rhythms, we removed the alpha frequency range using band-stop-filters (8-15 Hz) to avoid contributions from alpha rhythms (see Kosciessa, Kloosterman, & Garrett, 2020). A cluster-based permutation test indicated SampEn increases under probe uncertainty over posterior-occipital channels (Figure 5A). Notably, the magnitude of individual entropy modulation in this cluster scaled with increases in the SPMF [r(137) = 0.22, 95%CI [0.05, 0.37], p = 0.01], indicating that alpha desynchronization was accompanied by broadband changes in signal irregularity.

**Figure 5:**
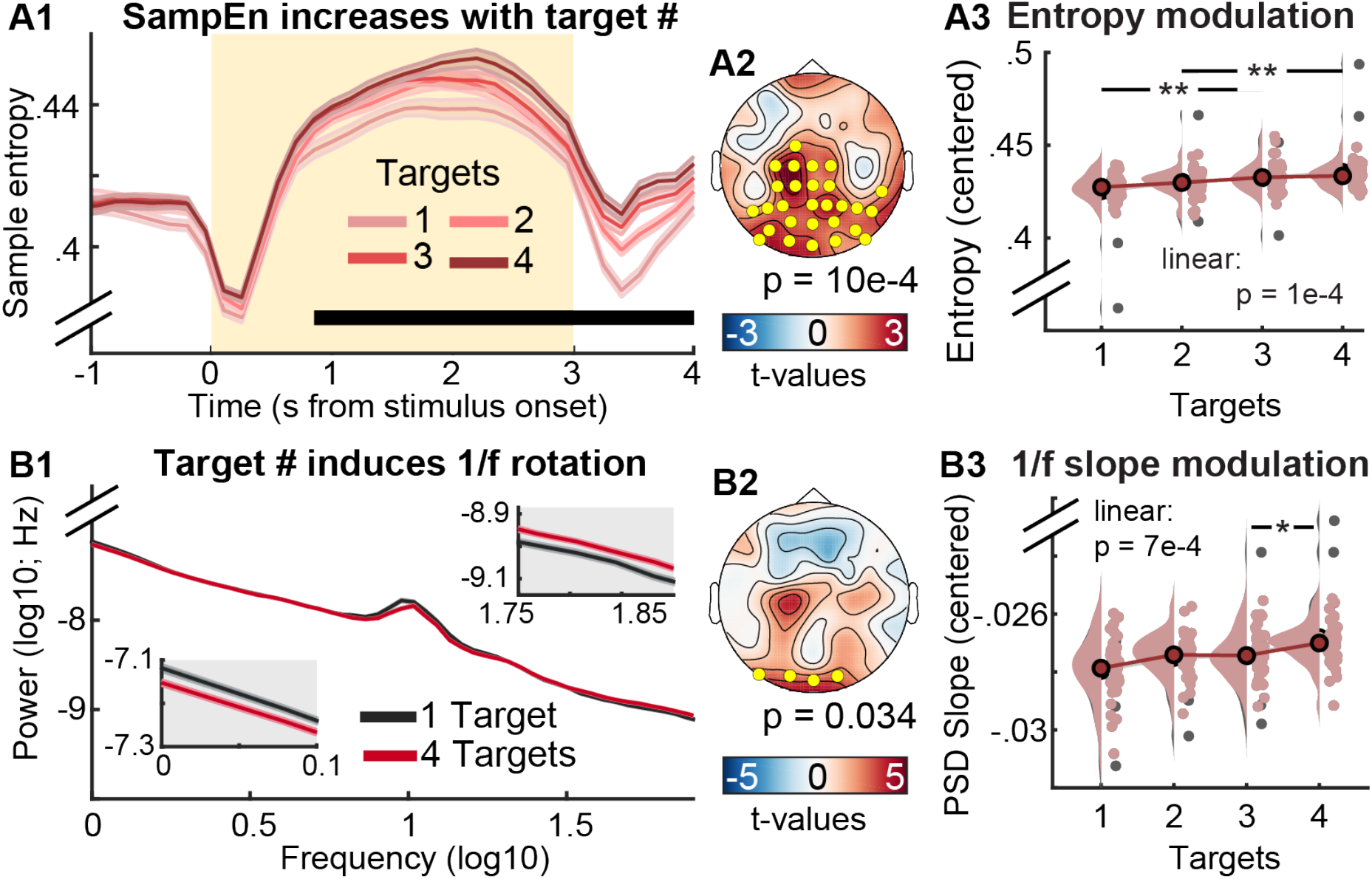
Uncertainty increases aperiodic dynamics during sensation as reflected in neural entropy (A) and 1/f slopes (B). (A1) Temporal traces of sample entropy (mean +− within-subject SE). The yellow background indicates the period of stimulus presentation. The black bar indicates time points at which permutation tests indicated linear load effects. (A2) Topography of linear load effect estimates, with yellow dots representing the significant cluster. (A3) Post-hoc analysis of entropy estimates within significant cluster. Grey dots indicate individual outliers (defined as Cook’s distance > 2.5*mean (Cook’s distance)) and have been removed from the statistical post-hoc assessment. Estimates have been within-subject centered for display purposes, while statistical analyses were run on uncentered data. (B1) Aperiodic slopes shallow with increased target load (i.e., spectral rotation across low- and high-frequencies; mean +− within-subject SE). Lower and upper insets highlight slope differences at low and high frequencies, respectively. (B2) Topography of linear load effects on 1/f slopes. Yellow dots indicate the significant occipital cluster used for post-hoc assessments. (B3) Same as A3, but for occipital aperiodic slopes. [*** p < .001, ** p < .01, * p < .05]

Aperiodic, scale-free spectral slopes are a major contributor to broadband SampEn, due to their joint sensitivity to autocorrelative structure (Kosciessa, Kloosterman, et al., 2020), and a shallowing of aperiodic (1/f) slopes has theoretically been associated with system excitability (Gao et al., 2017). We therefore assessed aperiodic slope changes during the stimulus period (excluding onset transients). In line with our hypothesis, participants’ PSD slopes shallowed under uncertainty (Figure 5B), suggesting that participants increased their excitatory tone in posterior cortex. In line with the expectation that sample entropy should be highly sensitive to scale-free dynamics, sample entropy was strongly related to individual PSD slopes across conditions (r = .77, p <.001) and with respect to linear changes in PSD slope with increasing uncertainty [r(137) = 0.44, 95%CI [0.3, 0.57], p = 4.92e-8]. In sum, heightened probe uncertainty desynchronized low-frequency alpha rhythms, and elevated the irregularity of cortical dynamics, in line with enhanced tonic excitability.

### Increases in phasic pupil diameter relate to transient excitability adjustments

Phasic arousal changes modulate perception and local cortical excitability (for reviews see Lee & Dan, 2012; McGinley, Vinck, et al., 2015). To test whether arousal increased alongside uncertainty, we assessed phasic changes in pupillometric responses as a proxy for arousal during stimulus presentation. We quantified phasic pupil responses via the 1^st^ temporal derivative (i.e. rate of change), as this measure has higher temporal precision and has been more strongly associated with noradrenergic responses than the overall pupil response (Reimer et al., 2014). Phasic pupil dilation systematically increased with probe uncertainty (Figure 6). This modulation occurred on top of a general pupil constriction due to stimulus-evoked changes in luminance (Figure 6A, inset), while the linear modulation occurred – by stimulus design – in the absence of systematic luminance changes.

**Figure 6:**
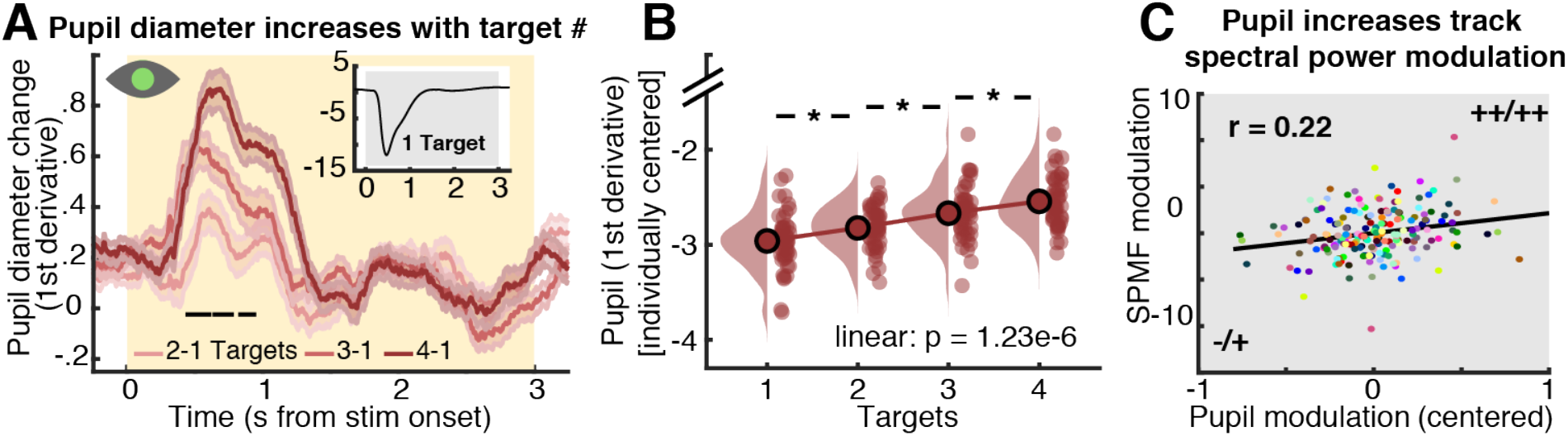
Effect of probe uncertainty on pupil diameter as a proxy for neuromodulation. (**A**) Phasic changes in pupil diameter increase with number of targets (mean +− within-subject SE). Significant linear load effects as indicated by a cluster-based permutation test are indicated via the black line. For follow-up analyses, we extracted median pupil values from 0 to 1.5 s. For display purposes but not statistics, derivative estimates were smoothed via application of a 200 ms median running average. (**B**) Post-hoc analysis of load effects in extracted median values. (**C**) Coupled changes between our spectral power modulation factor (SPMF) and pupil modulation. Dots represent linear model residuals (see methods), colored by participant. We indicate the direction of main effects for each variable via + and − (− = small decreases, −− = large decreases, + = small increases, ++ = large increases). [* p < .05]

Next, we assessed the relation between individual modulations in pupil diameter, cortical excitability and behavior. The magnitude of pupil increases tracked increases on the spectral power modulation factor (SPMF) [r(137) = 0.22, 95%CI [0.06, 0.38], p = 0.01], but did not directly relate to entropy [r(137) = −0.06, 95%CI [−0.23, 0.1], p = 0.45] or aperiodic slope changes [r(137) = −0.04, 95%CI [−0.2, 0.13], p = 0.67]. Participants with larger increases in pupil dilation also were faster integrators at baseline (r = .31, p = .033), and decreased integration more so with increasing probe uncertainty [r(137) = −0.17, 95%CI [−0.33, 0], p = 0.05], while showing more constrained NDT increases [r(137) = −0.21, 95%CI [−0.36, −0.04], p = 0.01]. This suggests that arousal jointly related to increases in local cortical excitability and subsequent choices.

### Thalamic BOLD modulation tracks excitability increases during sensation

Finally, we probed whether the thalamus acts as a subcortical nexus for sensory excitability adjustments under probe uncertainty. To allow spatially resolved insights into thalamic involvement, participants took part in a second, fMRI-based testing session during which they performed the same task. First, we investigated uncertainty-related changes in BOLD magnitude during stimulus processing via a task PLS. This analysis suggested two reliable (LV1: permuted *p* = .001; LV2: permuted *p* = .007) latent variables (Figure 7; see Table S1 for peak coordinates/statistics and Figure S5A, B for complete multivariate spatial patterns for the two LVs), with the first LV explaining the dominant amount of variance (89.6% crossblock covariance) compared to the second LV (8.7% crossblock covariance).

**Figure 7:**
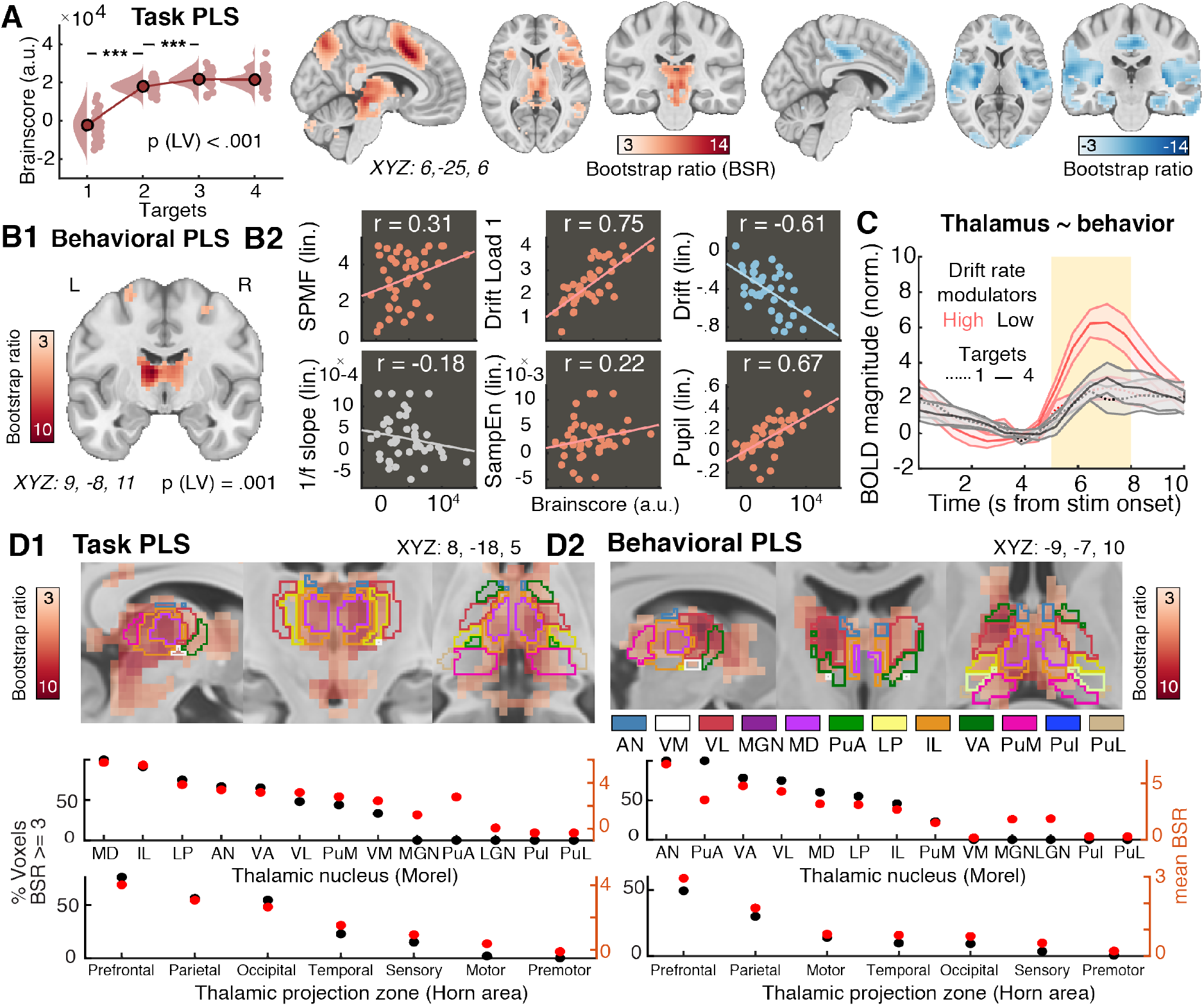
Upregulation of thalamic BOLD responses during stimulus processing is related to stronger excitability increases and better performance in upcoming decision task. (**A**) Results from multivariate task PLS investigating the relation of BOLD magnitude to attentional uncertainty. Data are individually centered across target loads. Activity maps show positive (left) and negative (right) bootstrap ratios of LV1, thresholded at a bootstrap ratio of 3 (p ~.001). Figure S5A presents the full loading matrices for LV1 and LV2. (**B**) Results from behavioral PLS, probing the association between linear changes in BOLD magnitude with behavioral, electrophysiological and pupillary changes under uncertainty. Figure S5B presents the complete factor loadings. (**C**) Visualization of thalamic modulation with uncertainty, split between low- and high-behavioral drift modulators (mean +− SE). The yellow shading indicates the approximate stimulus presentation period after accounting for the delay in the hemodynamic response function. Figure S5C plots all target conditions by group. (**D**) Thalamic expression pattern of the first task LV (D1) and the behavioral LV (D2). Scatters below indicate the major nuclei and projection zones in which behavioral relations are maximally reliable. For abbreviations see methods. Strongest expression is observed in antero-medial nuclei that project to fronto-parietal cortical targets. [*** p < .001]

The first latent variable (LV1) indicated load-related increases dominantly in cortical areas encompassing the fronto-parietal and the salience network, as well as thalamus. Primary positive contributors to LV1 (i.e., representing increases in BOLD with increasing probe uncertainty) were located in mid-cingulate cortex (MCG), inferior parietal lobule (IPL), bilateral anterior insula (aINS), inferior occipital gyrus (IOG), thalamus and bilateral inferior frontal gyrus (IFG). In contrast, relative uncertainty-related decreases in BOLD magnitude were dominantly observed in pallidum (potentially reflecting reduced motor preparation), bilateral posterior insula (pINS), left SFG, and left mid-cingulate cortex. Individual brain score increases were associated with stronger drift rate decreases [r(122) = −0.36, 95%CI [−0.5, −0.19], p = 5.11e-5], but not NDT, SPMF, or entropy (all p >.05). See Text S8 for results from the second latent variable (LV2), which might reflect decreased engagement at higher levels of target uncertainty.

Finally, we performed a behavioral PLS to probe whether regional BOLD modulation tracked a unified set of individual differences in the modulation of cortical excitability, arousal and behavior. In fact, we observed a single significant LV (permuted *p* = .001, 46.2% crossblock covariance) that dominantly loaded on anterior and midline thalamic nuclei with fronto-parietal projection zones (Figure 7D), and extended broadly across almost the entirety of thalamus. BOLD magnitude increases were more pronounced in participants exhibiting higher drift rates (i.e., more available evidence) (r = 0.75, 95% bootstrapped (bs) CI = [0.72,0.86]) and stronger drift reductions under probe uncertainty (r = −0.6, 95% bsCI = [−0.78,-0.54]; Figure 7B), as well as lower baseline non-decision times (r = −.37, 95% bsCI = [−.58, −.08]), confirming that increased thalamic responses reflected behaviorally adaptive contextual adjustments. This association was specific to the behavioral adjustments of interest, as we noted no relations with NDT modulation (r = .05, 95% bsCI = [−.31, .3]) or boundary separation (r = .08, 95% CI = [−.24, .37]). Importantly, higher (dominantly thalamic) BOLD modulation was further associated with greater increases on the SPMF (r = 0.31, 95% CI = [0.16,0.58]), in phasic pupil dilation (r = 0.67, 95% bsCI = [0.51,0.81]) and in entropy assessed during the EEG session (r = 0.22, 95% bsCI = [0.08,0.46]; Figure 7B). 1/f shallowing was not stably related to BOLD modulation (r = −0.18, 95% bsCI = [−0.38,0.17]), potentially due to noisier individual estimates. BOLD modulation was unrelated to chronological age (r = −.19, p = .21), gender (male vs. female; r = −.27, p = .08), subjective task difficulty (rated on 5-point Likert scale; r = −.02, p = .89), or framewise displacement of BOLD signals (an estimate of in-scanner motion; r = −.24, p = .13). Taken together, these results suggest a major role of the thalamus in integrating phasic neuromodulation to regulate rhythmic and aperiodic cortical excitability according to contextual demands.

## Discussion

To efficiently process information, cortical networks must be flexibly tuned to environmental demands. Invasive studies indicate a crucial role of the thalamus in such adaptations (for a review see Halassa & Kastner, 2017), but human evidence on thalamic involvement in rapid cortical regime switches at the service of behavioral flexibility has been missing. By combining a multi-modal experimental design with a close look at individual differences, we found that processing under contextual uncertainty is associated with a triad characterized by thalamic BOLD modulation, EEG-based cortical excitability, and pupil-based indicators of arousal. In the light of this triad, we propose that thalamic regulation of sensory excitability is crucial for adaptive sensory filtering in information-rich environments.

By cueing relevant dimensions of otherwise physically identical stimuli, we observed that increases in the number of attentional targets reliably reduced participants’ available evidence (as evidenced by drift rate decreases) during subsequent perceptual decisions. We interpret these changes as a negative (Dube, Emrich, & Al-Aidroos, 2017) but necessary and adaptive consequence of the need to encode multiple relevant features for an eventual decision regarding a single target. Concurrently, BOLD activity increased in the frontoparietal network (Dosenbach et al., 2007), composed of the inferior frontal junction (Zanto, Rubens, Thangavel, & Gazzaley, 2011), inferior frontal gyrus (Hampshire, Chamberlain, Monti, Duncan, & Owen, 2010), and posterior parietal cortex (Weerda, Vallines, Thomas, Rutschmann, & Greenlee, 2006; Wojciulik & Kanwisher, 1999), and the salience network (Uddin, 2015) – including anterior insula (Nelson et al., 2010) and dorsal anterior cingulate cortex (Weissman, Gopalakrishnan, Hazlett, & Woldorff, 2005). These cortical networks are thought to establish the contextual relevance of environmental stimuli, and to communicate this information to sensory cortex (Siegel et al., 2015). Accordingly, their BOLD activity often increases alongside multifaceted demands (see above), further in line with increased mediofrontal theta engagement (Cavanagh & Frank, 2014).

Besides such cortical responses at the group level however, we noted that individual increases in cortical excitability, drift rates, and arousal were tracked primarily by the extent of thalamic signal elevation, dominantly in areas with fronto-parietal projections. While past work emphasized the thalamic relay of peripheral information to cortex, recent theories highlight its dynamic involvement in cortical and cognitive function (for reviews see Dehghani & Wimmer, 2019; Halassa & Kastner, 2017; Halassa & Sherman, 2019; Pergola et al., 2018; Saalmann & Kastner, 2011; Ward, 2013; Wolff & Vann, 2019), with empirical support in humans (Garrett, Epp, Perry, & Lindenberger, 2018; Hwang, Bertolero, Liu, & D’Esposito, 2017; Shine et al., 2019), monkeys (Fiebelkorn, Pinsk, & Kastner, 2019; Saalmann, Pinsk, Wang, Li, & Kastner, 2012) and mice (Lewis et al., 2015; Schmitt et al., 2017; Wimmer et al., 2015). Notably, our task responds to demands for “tasks with multifaceted cognitive demands” (Pergola et al., 2018, p. 1017) to enhance sensitivity to higher-order thalamic involvement. In particular, anterior and midline thalamic nuclei, in which neuro-behavioral relations were maximal, may be essential for attentional set shifting (Marton, Seifikar, Luongo, Lee, & Sohal, 2018; Rikhye, Gilra, & Halassa, 2018; Wright, Vann, Aggleton, & Nelson, 2015) and to communicate such top-down information to sensory cortex via frontoparietal network coherence (Schmitt et al., 2017). Sensory processing in turn is shaped by thalamocortical transmission modes (Sherman, 2001). In ‘burst mode’, thalamic nuclei elicit synchronous activity that can boost stimulus detection (Alitto, Rathbun, Vandeleest, Alexander, & Usrey, 2019; Reinagel, Godwin, Sherman, & Koch, 1999) via non-linear gains of cortical responses (G. D. Smith, Cox, Sherman, & Rinzel, 2000; Swadlow & Gusev, 2001), whereas spike activity during ‘tonic mode’ more faithfully tracks incoming signals (Hartings, Temereanca, & Simons, 2003; Sherman, 2001). Shifts from sparse bursts towards tonic activity may underlie attention-related increases in thalamic BOLD magnitude observed here and in previous fMRI studies (Jagtap & Diwadkar, 2016; Kim, Cilles, Johnson, & Gold, 2012; Tomasi, Chang, Caparelli, & Ernst, 2007), although further work needs to elucidate the relation between thalamic transmission modes and BOLD responses (but see Liu et al., 2015).

Associated with thalamic bursting (Palva & Palva, 2007), cortical alpha rhythms may control sensory gain via periodic fluctuations in excitability (Dugue, Marque, & VanRullen, 2011; Haegens et al., 2011; Klimesch et al., 2007; Lorincz et al., 2009; Roux, Wibral, Singer, Aru, & Uhlhaas, 2013) that can signify rapid temporal imbalances between excitation and inhibition (Atallah & Scanziani, 2009; Poo & Isaacson, 2009). Supporting this notion, we observed a coupling between alpha phase and high-frequency power during stimulus processing, with participants engaging alpha rhythms most prevalently when prior cues afforded them a focus on single stimulus features (i.e., high available sensory evidence). Alpha rhythms have been consistently linked to the pulvinar nucleus (Halgren et al., 2019; Lopes da Silva, Vos, Mooibroek, & Van Rotterdam, 1980; Saalmann et al., 2012; Stitt, Zhou, Radtke-Schuller, & Frohlich, 2018), which also contributed to our multi-modal model. The pulvinar diffusely connects to visual and fronto-parietal cortices (Arcaro, Pinsk, & Kastner, 2015), affording it to build up contextual priors (Kanai, Komura, Shipp, & Friston, 2015; O’Reilly, Wyatte, & Rohrlich, 2017; Rikhye, Wimmer, & Halassa, 2018) that can regulate ‘bottom-up’ stimulus processing (Jaramillo, Mejias, & Wang, 2019), potentially via alpha rhythms (Saalmann et al., 2012; Suffczynski, Kalitzin, Pfurtscheller, & da Silva, 2001). While the localization of effects within the thalamus remains challenging in BOLD signals (Hwang et al., 2017), our results support a perspective in which alpha rhythms – shaped via thalamocortical circuits – dynamically extract relevant sensory information (Sadaghiani & Kleinschmidt, 2016) when contexts afford joint distractor suppression and target enhancement (Wöstmann, Alavash, & Obleser, 2019).

Complementing such selective gain control, overall increases in excitatory tone may serve multi-feature attention when only broad attentional guidance is available. Our results provide initial evidence that probe uncertainty transiently (a) desynchronizes alpha rhythms, (b) increases gamma power, and (c) elevates sample entropy while shallowing spectral slopes, a pattern that suggests increases in excitatory contributions to E/I mixture currents (Destexhe & Rudolph, 2004; Gao et al., 2017) and asynchronous neural firing (Destexhe et al., 2003). Conceptually, elevated excitability during high probe uncertainty facilitates an efficient and rapid switching between parallel feature activations. In agreement with this idea, joint activation of neural populations coding multiple relevant features has been observed during multi-feature attention (Mo et al., 2019). Furthermore, computational modeling indicates that E/I modulations in hierarchical networks optimally adjust multi-attribute choices (Pettine et al., 2020). Similar to our observation of enhanced excitability during probe uncertainty, Pettine et al. (2020) found increases in excitatory tone optimal for a linear weighting of multiple features, whereas inhibitory engagement increased the gain for specific features during more difficult perceptual decisions. As discussed above, such inhibitory tuning may regulate selective target gains via alpha rhythms, in line with the presumed importance of inhibitory interneurons in alpha rhythmogenesis (Lorincz et al., 2009).

Finally, probe uncertainty increased phasic pupil diameter, with strong links to parallel adjustments in behavior, EEG-based excitability, and thalamic BOLD modulation. Fluctuations in pupil diameter provide a non-invasive proxy of particularly noradrenergic drive in mice (Breton-Provencher & Sur, 2019; Reimer et al., 2014; Zerbi et al., 2019), monkeys (Aston-Jones & Cohen, 2005; Joshi, Li, Kalwani, & Gold, 2016) and humans (de Gee et al., 2017). As such, our results support neuromodulation as a potent regulator of excitability both directly at cortical targets (Constantinople & Bruno, 2011; McGinley, Vinck, et al., 2015) and via thalamic circuits (Liu et al., 2015; McCormick, 1989; McCormick, McGinley, & Salkoff, 2015; Schiff, 2008). Functionally, pupil diameter rises during states of heightened uncertainty (Krishnamurthy, Nassar, Sarode, & Gold, 2017; Nassar et al., 2012; Urai, Braun, & Donner, 2017), such as change points in dynamic environments (Murphy, Wilming, Hernandez-Bocanegra, Prat Ortega, & Donner, 2020; Nassar et al., 2012), and multi-feature attention (Alnaes et al., 2014; Koelewijn, Shinn-Cunningham, Zekveld, & Kramer, 2014), while increasing alongside cortical desynchronization (Dahl, Mather, Sander, & Werkle-Bergne, 2020; Murphy et al., 2020; Stitt et al., 2018; Waschke, Tune, & Obleser, 2019). Our results extend those observations, and suggest that neuromodulatory drive accompanies excitability increases especially when contexts prevent the formation of single attentional targets, potentially to serve a more faithful processing of complex environments (Berridge & Waterhouse, 2003; McGinley, David, et al., 2015).

Multiple neuromodulators, prominently noradrenaline and acetylcholine, regulate thalamocortical excitability (Lee & Dan, 2012; McCormick et al., 2015) and pupil responses (Reimer et al., 2014), but may differentially serve perceptual sensitivity vs. specificity demands (Shine, 2019). Specifically, noradrenergic drive may increase sensitivity to external stimuli (McCormick et al., 1991; Waterhouse & Navarra, 2019) by increasing E/I ratios (Froemke, Merzenich, & Schreiner, 2007; Martins & Froemke, 2015; Pfeffer et al., 2018), whereas cholinergic innervation might facilitate response selectivity (Bauer et al., 2012; Furey, Pietrini, & Haxby, 2000). However, as contrasting effects have also been observed for these modulators (e.g., Hirata, Aguilar, & Castro-Alamancos, 2006; Minces, Pinto, Dan, & Chiba, 2017; Vinck, Batista-Brito, Knoblich, & Cardin, 2015; Yu & Dayan, 2005), their functional separability necessitates further work.

To conclude, we report initial evidence that thalamocortical excitability adjustments guide human perception and decisions under uncertainty. Our results point to neuromodulatory changes regulated by the thalamus that trigger behaviorally relevant switches in cortical dynamics, from alpha-rhythmic gain control to increased tonic excitability, when contexts require a more faithful processing of information-rich environments. Given that difficulties in dealing with uncertainty, neuro-sensory hyperexcitability, and deficient E/I control are hallmarks of several clinical disorders (e.g., McFadyen, Dolan, & Garrido, 2020; Yang et al., 2016; Yizhar et al., 2011), we surmise that further research on individual differences in the modulation of contextual excitability might advance our understanding of cognitive flexibility in both healthy and diseased populations.

## Methods

### Sample

47 healthy young adults (18-35 years, mean age = 25.8 years, SD = 4.6, 25 women) performed a dynamic visual attention task during 64-channel active scalp EEG acquisition, 42 of whom returned for a subsequent 3T fMRI session. Due to participant and scanner availability, the average span between EEG and MR testing sessions was 9.8 days (SD = 9.5 days). Participants were recruited from the participant database of the Max Planck Institute for Human Development, Berlin, Germany (MPIB). Participants were right-handed, as assessed with a modified version of the Edinburgh Handedness Inventory (Oldfield, 1971), and had normal or corrected-to-normal vision. Participants reported to be in good health with no known history of neurological or psychiatric incidences, and were paid for their participation (10 € per hour). All participants gave written informed consent according to the institutional guidelines of the Deutsche Gesellschaft für Psychologie (DGPS) ethics board, which approved the study.

### Procedure: EEG Session

Participants were seated at a distance of 60 cm in front of a monitor in an acoustically and electrically shielded chamber with their heads placed on a chin rest. Following electrode placement, participants were instructed to rest with their eyes open and closed, each for 3 minutes. Afterwards, participants performed a standard Stroop task, followed by the visual attention task instruction & practice (see below), the performance of the task and a second Stroop assessment (Stroop results are not reported here). Stimuli were presented on a 60 Hz 1920×1080p LCD screen (AG Neovo X24) using PsychToolbox 3.0.11 (Brainard, 1997; Kleiner, Brainard, & Pelli, 2007; Pelli, 1997). The session lasted ~3 hours. EEG was continuously recorded from 60 active (Ag/AgCl) electrodes using BrainAmp amplifiers (Brain Products GmbH, Gilching, Germany). Scalp electrodes were arranged within an elastic cap (EASYCAP GmbH, Herrsching, Germany) according to the 10% system (Oostenveld & Praamstra, 2001), with the ground placed at AFz. To monitor eye movements, two additional electrodes were placed on the outer canthi (horizontal EOG) and one electrode below the left eye (vertical EOG). During recording, all electrodes were referenced to the right mastoid electrode, while the left mastoid electrode was recorded as an additional channel. Online, signals were digitized at a sampling rate of 1 kHz. In addition to EEG, we simultaneously tracked eye movements and assessed pupil diameter using EyeLink 1000+ hardware (SR Research, v.4.594) with a sampling rate of 1kHz.

### Procedure: MRI session

Forty-two participants returned for a second testing session that included structural and functional MRI assessments. First, participants took part in a short refresh of the visual attention task (“MAAT”, see below) instructions and practiced the task outside the scanner. Then, participants were located in the TimTrio 3T scanner and were instructed in the button mapping. We collected the following sequences: T1w, task (4 runs), T2w, resting state, DTI, with a 15 min out-of-scanner break following the task acquisition. The session lasted ~3 hours. Whole-brain task fMRI data (4 runs á ~11,5 mins, 1066 volumes per run) were collected via a 3T Siemens TrioTim MRI system (Erlangen, Germany) using a multi-band EPI sequence (factor 4; TR = 645 ms; TE = 30 ms; flip angle 60°; FoV = 222 mm; voxel size 3×3×3 mm; 40 transverse slices. The first 12 volumes (12 × 645 ms = 7.7 sec) were removed to ensure a steady state of tissue magnetization (total remaining volumes = 1054 per run). A T1-weighted structural scan was also acquired (MPRAGE: TR = 2500 ms; TE = 4.77 ms; flip angle 7°; FoV = 256 mm; voxel size 1×1×1 mm; 192 sagittal slices). A T2-weighted structural scan was also acquired (GRAPPA: TR = 3200 ms; TE = 347 ms; FoV = 256 mm; voxel size 1×1×1 mm; 176 sagittal slices).

### The multi-attribute attention task (“MAAT”)

We designed a task to parametrically control top-down attention to multiple feature dimensions, in the absence of systematic variation in bottom-up visual stimulation (see Figure 1). Participants attended a dynamic square display that jointly consisted of four attributes: color (red/green), movement direction (left, right), size (small, large) and saturation (low, high). The task incorporates features from random dot motion tasks which have been extensively studied in both animal models (Gold & Shadlen, 2007; Hanks & Summerfield, 2017; Siegel et al., 2015) and humans (Banca et al., 2015; Kelly & O’Connell, 2013). Following the presentation of these displays, a probe queried the prevalence of one of the four attributes in the display (e.g. whether the display comprised a greater proportion of either smaller or larger squares). Prior to stimulus onset, valid cue presentation informed participants about the active feature set, out of which one feature would be chosen as the probe. We parametrically manipulated uncertainty regarding the upcoming probe by systematically varying both the number and type of relevant features in the display.

The difficulty of each feature was determined by (a) the fundamental feature difference between the two alternatives and (b) the sensory evidence for each alternative in the display. For (a) the following values were used: high (RGB: 192, 255, 128) and low saturation green (RGB: 255, 128, 149) and high (RGB: 128, 255, 0) and low saturated red (RGB: 255, 0, 43) for color and saturation, 5 and 8 pixels for size differences and a coherence of .2 for directions. For (b) the proportion of winning to losing option (i.e. sensory evidence) was chosen as follows: color: 60/40; direction: 80/20; size: 65/35; luminance: 60/40. Parameter difficulty was established in a pilot population, with the aim to produce above-chance accuracy for individual features.

The experiment consisted of four runs of ~10 min, each consisting of eight blocks of eight trials (i.e., a total of 32 trial blocks; 256 trials). The size and constellation of the cue set was held constant within eight-trial blocks to reduce set switching and working memory demands. Each trial was structured as follows: cue onset during which the relevant targets were centrally presented (1 s), fixation phase (2 s), dynamic stimulus phase (3 s), probe phase (incl. response; 2 s); ITI (un-jittered; 1.5 s). At the onset of each block, the valid cue (attentional target set) was presented for 5 s. At the offset of each block, participants received feedback for 3 s. The four attributes spanned a constellation of 16 feature combinations (4×4), of which presentation frequency was matched within subjects. The size and type of cue set was pseudo-randomized, such that every size and constellation of the cue set was presented across blocks. Within each run of four blocks, every set size was presented once, but never directly following a block of the same set size. In every block, each feature in the active set acted as a probe in at least one trial. Moreover, any attribute equally often served as a probe across all blocks. Winning options for each feature were balanced across trials, such that (correct) button responses were equally distributed across the experiment. To retain high motivation during the task and encourage fast and accurate responses, we instructed participants that one response would randomly be drawn at the end of each block; if this response was correct and faster than the mean RT during the preceding block, they would earn a reward of 20 cents. However, we pseudo-randomized feedback such that all participants received a fixed payout of 10 € per session. This extra money was paid in addition to the participation fee at the end of the second session, at which point participants were debriefed.

### Behavioral estimates of probe-related decision processes

Sequential sampling models, such as the drift-diffusion model (DDM (Ratcliff & McKoon, 2008)), have been used to characterize evolving perceptual decisions in 2-alternative forced choice (2AFC) random dot motion tasks (Kelly & O’Connell, 2013), where the evolving decision relates to overt stimulus dynamics. In contrast with such applications, evidence integration here is tied to eidetic memory traces following the probe onset, similar to applications during memory retrieval (Ratcliff, 1978) or probabilistic decision making (Frank et al., 2015). Here, we estimated individual evidence integration parameters within the HDDM 0.6.0 toolbox (Wiecki, Sofer, & Frank, 2013) to profit from the large number of participants that can establish group priors for the relatively sparse within-subject data. Independent models were fit to data from the EEG and the fMRI session to allow reliability assessments of individual estimates. Premature responses faster than 250 ms were excluded prior to modeling, and the probability of outliers was set to 5%. 7000 Markov-Chain Monte Carlo samples were sampled to estimate parameters, with the first 5000 samples being discarded as burn-in to achieve convergence. We judged convergence for each model by visually assessing both Markov chain convergence and posterior predictive fits. Individual estimates were averaged across the remaining 2000 samples for follow-up analyses.

We fitted data to correct and incorrect RTs (termed ‘accuracy coding’ in Wiecki et al. (2013)). To explain differences in decision components, we compared four separate models. In the ‘full model’, we allowed the following parameters to vary between conditions: (i) the mean drift rate across trials, (ii) the threshold separation between the two decision bounds, (iii) the non-decision time, which represents the summed duration of sensory encoding and response execution. In the remaining models, we reduced model complexity, by only varying (a) drift, (b) drift + threshold, or (c) drift + NDT, with a null model fixing all three parameters. For model comparison, we first used the Deviance Information Criterion (DIC) to select the model which provided the best fit to our data. The DIC compares models on the basis of the maximal log-likelihood value, while penalizing model complexity. The full model provided the best fit to the empirical data based on the DIC index (Figure S1B) in both the EEG and the fMRI session. However, this model indicated an increase in decision thresholds (i.e., boundary separation) without an equivalent in the electrophysiological data (Figure S1C). We therefore fixed the threshold parameter across conditions, in line with previous work constraining model parameters on the basis of electrophysiological evidence (McGovern et al., 2018).

### EEG preprocessing

Preprocessing and analysis of EEG data were conducted with the FieldTrip toolbox (Oostenveld, Fries, Maris, & Schoffelen, 2011) and using custom-written MATLAB (The MathWorks Inc., Natick, MA, USA) code. Offline, EEG data were filtered using a 4^th^ order Butterworth filter with a pass-band of 0.5 to 100 Hz. Subsequently, data were down-sampled to 500 Hz and all channels were re-referenced to mathematically averaged mastoids. Blink, movement and heart-beat artifacts were identified using Independent Component Analysis (ICA; Bell & Sejnowski, 1995) and removed from the signal. Artifact-contaminated channels (determined across epochs) were automatically detected using (a) the FASTER algorithm (Nolan, Whelan, & Reilly, 2010), and by (b) detecting outliers exceeding three standard deviations of the kurtosis of the distribution of power values in each epoch within low (0.2-2 Hz) or high (30-100 Hz) frequency bands, respectively. Rejected channels were interpolated using spherical splines (Perrin, Pernier, Bertrand, & Echallier, 1989). Subsequently, noisy epochs were likewise excluded based on FASTER and on recursive outlier detection. Finally, recordings were segmented to participant cues to open their eyes, and were epoched into non-overlapping 3 second pseudo-trials. To enhance spatial specificity, scalp current density estimates were derived via 4^th^ order spherical splines (Perrin et al., 1989) using a standard 1005 channel layout (conductivity: 0.33 S/m; regularization: 1^-05; 14^th^ degree polynomials).

### Electrophysiological estimates of probe-related decision processes

#### Centroparietal Positive Potential (CPP)

The centroparietal positive potential (CPP) is an electrophysiological signature of internal evidence-to-bound accumulation (Kelly & O’Connell, 2013; McGovern et al., 2018; O’Connell et al., 2012). We probed the task modulation of this established signature and assessed its convergence with behavioral parameter estimates. To derive the CPP, preprocessed EEG data were low-pass filtered at 8 Hz with a 6^th^ order Butterworth filter to exclude low-frequency oscillations, epoched relative to response and averaged across trials within each condition. In accordance with the literature, this revealed a dipolar scalp potential that exhibited a positive peak over parietal channel POz (see Figure 2). We temporally normalized individual CPP estimates to a condition-specific baseline during the final 250 ms preceding probe onset. As a proxy of evidence drift rate, CPP slopes were estimates via linear regression from −250 ms to −100 ms surrounding response execution, while the average CPP amplitude from −50 ms to 50 ms served as an indicator of decision thresholds (i.e., boundary separation) (e.g., McGovern et al., 2018).

To investigate whether a similar ‘ramping’ potential was observed during stimulus presentation, we aligned data to stimulus onset and temporally normalized signals to the condition-specific signal during the final 250 ms prior to stimulus onset. During stimulus presentation, no ‘ramp’-like signal or load modulation was observed at the peak CPP channel. This suggests that immediate choice requirements were necessary for the emergence of the CPP, although prior work has shown the CPP to be independent of explicit motor requirements (O’Connell et al., 2012).

Finally, we assessed whether differences between probed stimulus attributes could account for load-related CPP changes (Figure S2C). For this analysis, we selected trials separately by condition and probed attribute. Note that for different probes, but not cues, trials were uniquely associated with each feature and trial counts were approximately matched across conditions. We explored differences between different conditions via paired t-tests. To assess load effects on CPP slopes and thresholds as a function of probed attribute, we calculated 1^st^-level load effects by means of a linear model, and assessed their difference from zero via paired t-tests.

#### Contralateral mu-beta

Decreases in contralateral mu-beta power provide a complementary, effector-specific signature of evidence integration (Donner et al., 2009; McGovern et al., 2018). We estimated mu-beta power using 7-cycle wavelets for the 8-25 Hz range with a step size of 50 ms. Spectral power was time-locked to probe presentation and response execution. We re-mapped channels to describe data recorded contra- and ipsi-lateral to the executed motor response in each trial, and averaged data from those channels to derive grand average mu-beta time courses. Individual average mu-beta time series were baseline-corrected using the −400 to −200 ms prior to probe onset, separately for each condition. For contralateral motor responses, remapped sites C3/5 and CP3/CP5 were selected based on the grand average topography for lateralized response executions (see inset in Figure S2A). As a proxy of evidence drift rate, mu-beta slopes were estimates via linear regression from −250 ms to −50 ms prior to response execution, while the average power −50 ms to 50 ms served as an indicator of decision thresholds (e.g., McGovern et al., 2018).

### Electrophysiological indices of top-down modulation during sensation

#### Low-frequency alpha and theta power

We estimated low-frequency power via a 7-cycle wavelet transform, using, for linearly spaced center frequencies in 1 Hz steps from 2 to 15 Hz. The step size of estimates was 50 ms, ranging from −1.5 s prior to cue onset to 3.5 s following stimulus offset. Estimates were log10-transformed at the single trial level (Smulders, ten Oever, Donkers, Quaedflieg, & van de Ven, 2018), with no explicit baseline.

#### High-frequency gamma power

Gamma responses were estimated using multi-tapers (five tapers; discrete prolate spheroidal sequences) with a step size of 200 ms, a window length of 400 ms and a frequency resolution of 2.5 Hz. The frequency range covered frequencies between 45-90 Hz, with spectral smoothing of 8 Hz. Estimates were log10-transformed at the single trial level. We normalized individual gamma-band responses via single-trial z-normalization. In particular, for each frequency, we subtracted single-trial power −700 to −100 ms prior to stimulus onset, and divided by the standard deviation of power values during the same period. Finally, to account for baseline shifts during the pre-stimulus period, we subtracted condition-wise averages during the same baseline period.

#### Multivariate assessment of spectral power changes with stimulus onset and uncertainty

To determine changes in spectral power upon stimulus onset, and during stimulus presentation with load, we entered individual power values into multivariate partial least squares (PLS) analyses (see *Multivariate partial least squares analyses*) using the MEG-PLS toolbox [version 2.02b] (Cheung, Kovacevic, Fatima, Misic, & McIntosh, 2016). We concatenated low- (2-15 Hz) and high-frequency (45-90 Hz) power matrices to assess joint changes in the PLS models. To examine a multivariate contrast of spectral changes upon stimulus onset (averaged across conditions) with spectral power in the pre-stimulus baseline period, we performed a task PLS on data ranging from 500 ms pre-stim to 500 ms post-stim. Temporal averages from −700 to −100 ms pre-stimulus onset were subtracted as a baseline. To assess power changes as a function of probe uncertainty, we segmented the data from 500 ms post stim onset to stimulus offset (to exclude transient evoked onset responses), and calculated a task PLS concerning the relation between experimental uncertainty conditions and time-space-frequency power values. As a control, we performed a behavioral PLS analysis to assess the relevance of individual frequency contributions to the behavioral relation. For this analysis, we computed linear slopes (target amount) for each time-frequency point at the 1^st^ (within-subject) level, which were subsequently entered into the 2^nd^ level PLS analysis. On the behavioral side, we assessed both linear changes in pupil diameter, as well as drift rates during selective attention and linear decreases in drift rate under uncertainty. Finally, spontaneous fluctuations in pre-stimulus power have been linked to fluctuations in cortical excitability (Iemi, Chaumon, Crouzet, & Busch, 2017; Lange, Oostenveld, & Fries, 2013). We thus probed the role of upcoming processing requirements on pre-stimulus oscillations, as well as the potential relation to behavioral outcomes using task and behavioral PLS analyses. The analysis was performed as described above, but restricted to time points occurring during the final second prior to stimulus onset.

#### Steady State Visual Evoked Potential (SSVEP)

The SSVEP characterizes the phase-locked, entrained visual activity (here 30 Hz) during dynamic stimulus updates (e.g., Ding, Sperling, & Srinivasan, 2006). These features differentiate it from induced broadband activity or muscle artefacts in similar frequency bands. We used these properties to normalize individual single-trial SSVEP responses prior to averaging: (a) we calculated an FFT for overlapping one second epochs with a step size of 100 ms (Hanning-based multitaper), averaged them within each load condition and, (b) spectrally normalized 30 Hz estimates by subtracting the average of estimates at 28 and 32 Hz, effectively removing broadband effects (i.e., aperiodic slopes), (c) and finally, we subtracted a temporal baseline −700 to −100 ms prior to stimulus onset. Linear load effects on SSVEPs were assessed by univariate cluster-based permutation tests on channel x time data (see *Univariate statistical analyses using cluster-based permutation tests*).

#### Time-resolved sample entropy

Sample entropy (Richman & Moorman, 2000) quantifies the irregularity of a time series of length *N* by assessing the conditional probability that two sequences of *m* consecutive data points will remain similar when another sample (*m+1*) is included in the sequence (for a visual example see Figure 1A). Sample entropy is defined as the inverse natural logarithm of this conditional similarity: 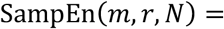 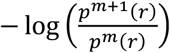. The similarity criterion (*r*) defines the tolerance within which two points are considered similar and is defined relative to the standard deviation (~variance) of the signal (here set to r = .5). We set the sequence length *m* to 2, in line with previous applications (Kosciessa, Kloosterman, et al., 2020). An adapted version of sample entropy calculations was used (Grandy, Garrett, Schmiedek, & Werkle-Bergner, 2016; Kloosterman, Kosciessa, Lindenberger, Fahrenfort, & Garrett, 2019; Kosciessa, Kloosterman, et al., 2020), wherein entropy is estimated across discontinuous data segments to provide time-resolved estimates. The estimation of scale-wise entropy across trials allows for an estimation of coarse scale entropy also for short time-bins, i.e., without requiring long, continuous signals, while quickly converging with entropy estimates from continuous recordings (Grandy et al., 2016). To remove the influence of posterior-occipital low-frequency rhythms on entropy estimates, we notch-filtered the 8-15 Hz alpha band using 6^th^ order Butterworth filter prior to the entropy calculation (Kosciessa, Kloosterman, et al., 2020). Time-resolved entropy estimates were calculated for 500 ms windows from −1 s pre-stimulus to 1.25 s post-probe with a step size of 150 ms. As entropy values are implicitly normalized by the variance in each time bin via the similarity criterion, no temporal baselining was used. Linear load effects on entropy were assessed by univariate cluster-based permutation tests on channel x time data (see *Univariate statistical analyses using cluster-based permutation tests*).

#### Aperiodic (1/f) slopes

The aperiodic 1/f slope of neural recordings is closely related to the sample entropy of broadband signals (Kosciessa, Kloosterman, et al., 2020), and has been suggested as a proxy for ‘cortical excitability’ and excitation-inhibition balance (Gao et al., 2017). Spectral estimates were computed by means of a Fast Fourier Transform (FFT) over the final 2.5 s of the presentation period (to exclude onset transients) for 41 logarithmically spaced frequencies between 2 and 64 Hz (Hanning-tapered segments zero-padded to 10 s) and subsequently averaged. Spectral power was log10-transformed to render power values more normally distributed across subjects. Power spectral density (PSD) slopes were derived by linearly regressing log-transformed power values on log-transformed frequencies. The spectral range from 7-13 Hz was excluded from the background fit to exclude a bias by the narrowband alpha peak (Kosciessa, Kloosterman, et al., 2020) and thus to increase the specificity to aperiodic variance. Linear load effects on 1/f slopes were assessed by univariate cluster-based permutation tests on channel data (see *Univariate statistical analyses using cluster-based permutation tests*).

#### Rhythm-specific estimates

Spectral power estimates conflate rhythmicity with aperiodic events in time, space and magnitude (Kosciessa, Grandy, Garrett, & Werkle-Bergner, 2020). Given that we observed changes in aperiodic slopes, we verified that observed narrowband effects in the theta and alpha band describe narrowband changes in rhythmicity. For this purpose, we identified single-trial spectral events using the extended BOSC method (Caplan, Madsen, Raghavachari, & Kahana, 2001; Kosciessa, Grandy, et al., 2020; Whitten, Hughes, Dickson, & Caplan, 2011). In short, this method identifies stereotypic ‘rhythmic’ events at the single-trial level, with the assumption that such events have significantly higher power than the 1/f background and occur for a minimum number of cycles at a particular frequency. This procedure dissociates narrowband spectral peaks from the aperiodic background spectrum. Here, we used a three-cycle threshold during detection, while defining the power threshold as the 95^th^ percentile above the individual background power. A 5-cycle wavelet was used to provide the time-frequency transformations for 49 logarithmically-spaced center frequencies between 1 and 64 Hz. Rhythmic episodes were detected as described in (Kosciessa, Grandy, et al., 2020). Prior to fitting the 1/f slopes, the most dominant individual rhythmic alpha peak between 8 and 15 Hz was removed, as well as the 28-32 Hz range, to exclude the SSVEP. Detection of episodes was restricted to the time of stimulus presentation, excluding the first 500 ms to reduce residual pre-stimulus activity and onset transients. Within each participant and channel, the duration and SNR of individual episodes with a mean frequency between 4-8 Hz (Theta) and 8-15 Hz (Alpha) were averaged across trials. Effects of target number were assessed within the averaged spatial clusters indicated in Figure 3 by means of paired t-tests.

### Alpha-gamma phase-amplitude coupling (PAC)

We assessed alpha-phase-to-gamma-amplitude coupling to assess the extent of phasic modulation of gamma power within the alpha band. As phase information is only interpretable during the presence of a narrowband rhythm (Aru et al., 2015), we focused our main analysis on 250 ms time segments following the estimated onset of a rhythm in the 8-15 Hz alpha range (see *Rhythm-specific estimates* above; Figure 4A). This time window ensured that segments fulfilled the 3-cycle criterion imposed during eBOSC rhythm detection to ensure that a rhythm was present. We selected three occipital channels with maximal gamma power (O1, O2, Oz; shown in Figure 4A) and pooled detected alpha episodes across these channels. We pooled data across load conditions, as we observed no consistent PAC within individual load conditions (data not shown), perhaps due to low episode counts. To derive the alpha carrier phase, we band-pass filtered signals in the 8-15 Hz band, and estimated the analytic phase time series via Hilbert transform. For the amplitude of modulated frequencies, we equally applied band-pass filters from 40 to 150 Hz (step size: 2 Hz), with adaptive bandwidths (+/− 20% of center frequency). Filtering was implemented using MATLAB’s acausal filtfilt() routine using linear finite impulse response (FIR) filters with an adaptive filter order set as 3 times the ratio of the sampling frequency to the low-frequency cutoff (Tort et al., 2008). For each applied bandpass filter, we removed 250 ms at each edge to avoid filter artifacts. For each frequency, narrowband signals were z-scored to normalize amplitudes across frequencies, and absolute values of the Hilbert-derived complex signal were squared to produce instantaneous power time series. We estimated the MI between the 8-15 Hz phase and high-frequency power via normalized entropy (Tort et al., 2008) using 16 phase bins. Power estimates were normalized by dividing the bin-specific power by the sum of power across bins. To make MI estimates robust against random coupling, we estimated MI for 1000 surrogate data, which shuffled the trial association of phase and amplitude information. We subtracted the mean surrogate MI value from the original MI index for a final, surrogate-normalized MI estimate. The resulting MI estimates across frequencies were then subjected to a cluster-based permutation test to assess significant clusters from zero using paired t-tests. For Figure 4B, we followed the procedure by Canolty et al. (2006). Alpha troughs were identified as local minima of phases < [−pi+.01]. For visualization, data were averaged across center frequencies from 80-150 Hz, as significant coupling overlapped with this range. We performed identical analyses for the 250 ms periods prior to rhythm onset (grey shading in Figure 4A) as a control condition. We performed analogous phase-amplitude-coupling analyses for the Mean Vector Length (MVL; Canolty et al., 2006) index, with comparable results (data not shown).

### Analyses of pupil diameter

Pupil diameter was recorded during the EEG session using EyeLink 1000 at a sampling rate of 1000 Hz, and was analyzed using FieldTrip and custom-written MATLAB scripts. Blinks were automatically indicated by the EyeLink software (version 4.40). To increase the sensitivity to periods of partially occluded pupils or eye movements, the first derivative of eye-tracker-based vertical eye movements was calculated, z-standardized and outliers >= 3 STD were removed. We additionally removed data within 150 ms preceding or following indicated outliers. Finally, missing data were linearly interpolated and data were epoched to 3.5 s prior to stimulus onset to 1 s following stimulus offset. We quantified phasic arousal responses via the 1^st^ temporal derivative (i.e. rate of change) of pupil diameter traces, as this measure (i) has higher temporal precision and (ii) has been more strongly associated with noradrenergic responses than the overall response (Reimer et al., 2014). We downsampled pupil time series to 200 Hz. For visualization, but not statistics, we smoothed pupil traces using a moving average median of 200 ms. We statistically assessed a linear load effect using a cluster-based permutation test on the 1D pupil traces (see *Univariate statistical analyses using cluster-based permutation tests*). For post-hoc assessments, we extracted the median pupil derivative during the first 1.5 s following stimulus onset.

### fMRI-based analyses

#### Preprocessing of functional MRI data

fMRI data were preprocessed with FSL 5 (RRID:SCR_002823) (Jenkinson, Beckmann, Behrens, Woolrich, & Smith, 2012; S. M. Smith et al., 2004). Pre-processing included motion correction using McFLIRT, smoothing (7mm) and high-pass filtering (.01 Hz) using an 8^th^ order zero-phase Butterworth filter applied using MATLAB’s filtfilt function. We registered individual functional runs to the individual, ANTs brain-extracted T2w images (6 DOF), to T1w images (6 DOF) and finally to 3mm standard space (ICBM 2009c MNI152 nonlinear symmetric) (Fonov et al., 2011) using nonlinear transformations in ANTs (Avants et al., 2011). (For one participant, no T2w image was acquired and 6 DOF transformation of BOLD data was preformed directly to the T1w structural scan.) We then masked the functional data with the ICBM 2009c GM tissue prior (thresholded at a probability of 0.25), and detrended the functional images (up to a cubic trend) using SPM8.

We also used a series of extended preprocessing steps to further reduce potential non-neural artifacts (Garrett, Kovacevic, McIntosh, & Grady, 2010; Garrett et al., 2015). Specifically, we examined data within-subject, within-run via spatial independent component analysis (ICA) as implemented in FSL-MELODIC (Beckmann & Smith, 2004). Due to the high multiband data dimensionality in the absence of low-pass filtering, we constrained the solution to 30 components per participant. Noise components were identified according to several key criteria: a) Spiking (components dominated by abrupt time series spikes); b) Motion (prominent edge or “ringing” effects, sometimes [but not always] accompanied by large time series spikes); c) Susceptibility and flow artifacts (prominent air-tissue boundary or sinus activation; typically represents cardio/respiratory effects); d) White matter (WM) and ventricle activation (Birn, 2012); e) Low-frequency signal drift (A. M. Smith et al., 1999); f) High power in high-frequency ranges unlikely to represent neural activity (≥ 75% of total spectral power present above .10 Hz;); and g) Spatial distribution (“spotty” or “speckled” spatial pattern that appears scattered randomly across ≥ 25% of the brain, with few if any clusters with ≥ 80 contiguous voxels [at 2×2×2 mm voxel size]). Examples of these various components we typically deem to be noise can be found in (Garrett, McIntosh, & Grady, 2014). By default, we utilized a conservative set of rejection criteria; if manual classification decisions were challenging due to mixing of “signal” and “noise” in a single component, we generally elected to keep such components. Three independent raters of noise components were utilized; > 90% inter-rater reliability was required on separate data before denoising decisions were made on the current data. Components identified as artifacts were then regressed from corresponding fMRI runs using the regfilt command in FSL.

To reduce the influence of motion and physiological fluctuations, we regressed FSL’s 6 DOF motion parameters from the data, in addition to average signal within white matter and CSF masks. Masks were created using 95% tissue probability thresholds to create conservative masks. Data and regressors were demeaned and linearly detrended prior to multiple linear regression for each run. To further reduce the impact of potential motion outliers, we censored significant DVARS outliers during the regression as described by (Power et al., 2014). In particular, we calculated the ‘practical significance’ of DVARS estimates and applied a threshold of 5 (Afyouni & Nichols, 2018). The regression-based residuals were subsequently spectrally interpolated during DVARS outliers as described in (Power et al., 2014) and (Parkes, Fulcher, Yucel, & Fornito, 2018). BOLD analyses were restricted to participants with both EEG and MRI data available (N = 42).

#### 1^st^ level analysis: univariate beta weights for load conditions

We conducted a 1^st^ level analysis using SPM12 to identify beta weights for each load condition separately. Design variables included stimulus presentation by load (4 volumes; parametrically modulated by sequence position), onset cue (no mod.), probe (2 volumes, parametric modulation by RT).

Design variables were convolved with a canonical HRF, including its temporal derivative as a nuisance term. Nuisance regressors included 24 motion parameters (Friston, Williams, Howard, Frackowiak, & Turner, 1996), as well as continuous DVARS estimates. Autoregressive modelling was implemented via FAST. Output beta images for each load condition were finally averaged across runs.

#### 2^nd^ level analysis: Multivariate modulation of BOLD responses

We investigated the multivariate modulation of the BOLD response at the 2^nd^ level using PLS analyses (see *Multivariate partial least squares analyses*). Specifically, we probed the relationship between voxel-wise 1^st^ level beta weights and probe uncertainty within a task PLS. Next, we assessed the relationship between task-related BOLD signal changes and interindividual differences in the joint modulation of decision processes, cortical excitability, and pupil modulation by means of a behavioral PLS. For this, we first calculated linear slope coefficients for voxel-wise beta estimates. Then, we included behavioral variables including HDDM parameter estimates during selective attention, as well as linear changes with load, individual linear condition modulation of the following variables: multivariate spectral power, pupil dilation, 1/f modulation and entropy residuals. Prior to these covariates in the model, we visually assessed whether the distribution of linear changes variables was approximately Gaussian. In the case of outliers (as observed for the SPMF, 1/f slopes, and entropy), we winsorized values at the 95^th^ percentile. For visualization, spatial clusters were defined based on a minimum distance of 10 mm, and by exceeding a size of 25 voxels. We identified regions associated with peak activity based on cytoarchitectonic probabilistic maps implemented in the SPM Anatomy Toolbox (Version 2.2c) (Eickhoff et al., 2005). If no assignment was found, the most proximal assignment to the coordinates reported in Table S1 within the cluster was reported.

#### Temporal dynamics of thalamic engagement

To visualize the modulation of thalamic activity by load, we extracted signals within a binary thalamic mask extracted from the Morel atlas, including all subdivisions. Preprocessed BOLD timeseries were segmented into trials, spanning the period from the stimulus onset to the onset of the feedback phase. Given a time-to-peak of a canonical hemodynamic response function (HRF) between 5-6 seconds, we designated the 3 second interval from 5-8 seconds following the stimulus onset trigger as the stimulus presentation interval, and the 2 second interval from 3-5 s as the fixation interval, respectively. Single-trial time series were then temporally normalized to the temporal average during the approximate fixation interval. To visualize inter-individual differences in thalamic engagement, we performed a median split across participants based on their individual drift modulation.

#### Thalamic loci of behavioral PLS

To assess the thalamic loci of most reliable behavioral relations (Figure S5C), we assessed bootstrap ratios within two thalamic masks. First, for nucleic subdivisions, we used the Morel parcellation scheme as consolidated and kindly provided by (Hwang et al., 2017) for 3 mm data at 3T field strength. The abbreviations are as follows: AN: anterior nucleus; VM: ventromedial; VL: ventrolateral; MGN: medial geniculate nucleus; LGN: lateral geniculate nucleus; MD: mediodorsal; PuA: anterior pulvinar; LP: lateral-posterior; IL: intra-laminar; VA: ventral-anterior; PuM: medial pulvinar; Pul: pulvinar proper; PuL: lateral pulvinar. Second, to assess cortical white-matter projections we considered the overlap with seven structurally-derived cortical projection zones suggested by (Horn & Blankenburg, 2016), which were derived from a large adult sample (*N* = 169). We binarized continuous probability maps at a relative 75% threshold of the respective maximum probability, and re-sliced masks to 3 mm size.

### Statistical analyses

#### Assessment of covarying load effect magnitudes between measures

To assess a linear modulation of dependent variables, we calculated 1^st^ level beta estimates for the effect of load (y = intercept+β*LOAD+e) and assessed the slope difference from zero at the group level using paired t-tests. We assessed the relation of individual load effects between measures of interest by means of partial repeated measures correlations. In a simplified form, repeated measured correlation (Bakdash & Marusich, 2017) fits a linear model between two variables x1 and x2 of interest, while controlling for repeated assessments within subjects [x1~1+β1*ID+β2*x2+e] (1). Crucially, to exclude bivariate relations that exclusively arise from joint main effects of number of targets, we added target load as an additional categorical covariate [x1~1+β1*ID+β2*LOAD+β3*x2+e] (2) to remove group condition means. Resulting estimates characterize the group-wise coupling in the (zero-centered) magnitude of changes between the DV and the IV across the four load levels. To identify the directionality of the coupling, we assessed the direction of main effects for x1 and x2. We statistically compared this model to a null model without the term of interest [x1~1+β1*ID+β2*LOAD +e] (3) to assess statistical significance. We report the bivariate residual effect size by assessing the square root of partial eta squared. We extend this model with additional beta*covariate terms when reporting control for additional covariates.

#### Within-subject centering

To better visualize effects within participants, we use within-subject centering across repeated measures conditions by subtracting individual condition means, and adding global means. For these visualizations, only the mean of the dependent values is directly informative, as the plotted spread reflects within-subject, and not between-subject, variation. This procedure is similar to the creation of within-subject standard errors. Within-subject centering is exclusively used for display, but not statistical calculations.

#### Univariate cluster-based permutation analyses

For data with a low-dimensional structure (e.g., based on a priori averaging or spatial cluster assumptions), we used univariate cluster-based permutation analyses (CBPAs) to assess significant modulations by target load or with stimulus onset. These univariate tests were performed by means of dependent samples t-tests; cluster-based permutation tests (Maris & Oostenveld, 2007) were performed to control for multiple comparisons. Initially, a clustering algorithm formed clusters based on significant t-tests of individual data points (p <.05, two-sided; cluster entry threshold) with the spatial constraint of a cluster covering a minimum of three neighboring channels. Then, the significance of the observed cluster-level statistic, based on the summed t-values within the cluster, was assessed by comparison to the distribution of all permutation-based cluster-level statistics. The final cluster p-value that we report in all figures was assessed as the proportion of 1000 Monte Carlo iterations in which the cluster-level statistic was exceeded. Cluster significance was indicated by p-values below .025 (two-sided cluster significance threshold).

#### Multivariate partial least squares analyses

For data with a high-dimensional structure, we performed multivariate partial least squares analyses (Krishnan, Williams, McIntosh, & Abdi, 2011; McIntosh, Bookstein, Haxby, & Grady, 1996; McIntosh & Lobaugh, 2004). To assess main effect of probe uncertainty or stimulus onset, we performed Task PLS analyses. Task PLS begins by calculating a between-subject covariance matrix (COV) between conditions and each neural value (e.g., time-space-frequency power), which is then decomposed using singular value decomposition (SVD). This yields a left singular vector of experimental condition weights (U), a right singular vector of brain weights (V), and a diagonal matrix of singular values (S). Task PLS produces orthogonal latent variables (LVs) that reflect optimal relations between experimental conditions and the neural data. To examine multivariate relations between neural data and other variables of interest, we performed behavioral PLS analyses. This analysis initially calculates a between-subject correlation matrix (CORR) between (1) each brain index of interest (e.g., spectral power, 1^st^ level BOLD beta values) and (2) a second ‘behavioral’ variable of interest (note that although called behavioral, this variable can reflect any variable of interest, e.g., behavior, pupil dilation, spectral power). CORR is then decomposed using singular value decomposition (SVD): SVD_CORR_ = *USV’*, which produces a matrix of left singular vectors of cognition weights (*U*), a matrix of right singular vectors of brain weights (*V*), and a diagonal matrix of singular values (*S*). For each LV (ordered strongest to weakest in *S*), a data pattern results which depicts the strongest available relation to the variable of interest. Significance of detected relations of both PLS model types was assessed using 1000 permutation tests of the singular value corresponding to the LV. A subsequent bootstrapping procedure indicated the robustness of within-LV neural saliences across 1000 resamples of the data (Efron & Tibshirani, 1986). By dividing each brain weight (from *V*) by its bootstrapped standard error, we obtained “bootstrap ratios” (BSRs) as normalized robustness estimates. We generally thresholded BSRs at values of ±3.00 (∼99.9% confidence interval). We also obtained a summary measure of each participant’s robust expression of a particular LV’s pattern (a within-person “brain score”) by either (1) multiplying the vector of brain weights (*V)* from each LV by within-subject vectors of the neural values (separately for each condition within person) for the Task PLS models, or (2) in the behavioral PLS model, by multiplying the model-based vector of weights (*V*) by each participant’s vector of neural values (*P*), producing a single within-subject value: Brain score = *VP’*.

## Acknowledgements

We thank our research assistants and participants for their contributions to the present work, Alistair Perry for assistance in fMRI preprocessing, and Steffen Wiegert for organizational support.

## Author contributions

JQK: Conceptualization, Methodology, Investigation, Software, Formal analysis, Visualization, Writing – original draft, Writing – review and editing, Validation, Data Curation; UL: Conceptualization, Resources, Writing – review and editing, Supervision, Funding acquisition; DDG: Conceptualization, Methodology, Software, Resources, Writing—review and editing, Supervision, Project administration, Funding acquisition.

## Data and Code Availability

Experiment code is available from https://git.mpib-berlin.mpg.de/LNDG/multi-attribute-task. Primary EEG and fMRI data (excluding structural images exempt from informed consent) will be made available following publication. Code to reproduce the analyses will be made available at https://git.mpib-berlin.mpg.de/LNDG/stateswitch.

## Text S1. Parameter interrelations

To better understand individual differences in behavioral performance, we explored inter-individual associations between model parameter estimates and ‘raw’ median RT and mean accuracy. Linear drift rate decreases were inter-individually associated with decreases in accuracy (EEG: r = .35, p = .015, MRI: r = .46, p = .001), but not RT increases (both p > .05), whereas non-decision-time (NDT) increases tracked individual RT increases (EEG: r = .56, p = 3e-5, MRI: r = .64, p = 2e-6), but not accuracy decreases (both p > .05). For single targets, faster RTs were associated with larger drift rates (EEG: r = −.63, p = 3e-6, MRI: r = −.47, p = .002), lower non-decision times (EEG: r = .41, p = .005, MRI: r = .58, p = 3e-5), and lower boundary separation (EEG: r = .58, p = 3e-5, MRI: r = .5, p = 6e-4). More accurate performance for single targets was related to higher drift rates (EEG: r = .74, p = 3e-9; MRI: r = .79, p = 3e-10), but unrelated to boundary separation (EEG: r = .23, p = .121, MRI: r = .18, p = .244) or non-decision times (EEG: r = −.27, p = .069, MRI: r = −.38, p = .011). Amongst model parameters, we observed no parameter relations for single targets (all p > .05). However, we observed intercept-change correlations: subjects with larger drift rates for single targets exhibited strong linear drift rate reductions (EEG: r = −.93, p = 4e-22, MRI: r = −.88, p = 1e-15). Moreover, subjects with larger boundary separation showed stronger linear increases in non-decision time (r = .46, p = 9e-4, MRI: r = .59, p =2e-5). Non-decision time under selective attention, putatively dominantly reflecting visual encoding time, did not relate to changes in drift rate or NDT (both p > .05). Similarly, boundary separation did not relate to drift rate decreases (both p > .05) and drift rates under selective attention were unrelated to NDT increases (both p > .05).

## Text S2. Behavioral benefits due to convergent responses

To reduce response mapping demands following probe presentation, we fixed response mapping for the two options of each feature throughout the experiment. Given that multiple attributes converge onto a similar response in a given trial, the potential to prepare motor action prior to probe presentation co-varies as a function of load. To assess the influence of this response agreement on our results, we ran an additional HDDM that simultaneously modelled both a main effect of load, as well as categorical response agreement. Notably, the obtained target load effects on drift rate and NDT were virtually identical to those observed in the selected model in both sessions (reliability of all linear effects: r >= .9 p <.001; data not shown), while linear decreases in drift and increases in NDT were also observed as a function of response divergence (i.e., lower drift and higher NDT if the probed attribute required a differential response than the other cued attributes; shown in Figure S1D for the EEG session; qualitatively similar results were obtained for MRI session; all linear effects p < .001). This suggests that response agreement systematically impacted decision processes, but cannot account for the main effects of target load. However, the large amount of added model parameters introduced partial convergence issues. We therefore chose the simpler model without response agreement for our main analyses.

## Text S3. NDT increases indicate extended motor preparation demands

We observed a parametric increase in non-decision time (NDT) with target uncertainty (Figure 2B) that described shifts in RT distribution onset (Figure S3A). NDT is thought to characterize the duration of processes preceding and following evidence accumulation, i.e., probe encoding and planning/execution of the motor response. We therefore examined sensory probe- and response-related ERP components regarding their modulation by prior target uncertainty. We time-locked the CPP to the NDT group estimate for a single target – for which no button remapping was required – and (2) to the condition-wise NDT estimate. However, we observed no shift in CPP onset (Figure S3B), suggesting constant visual encoding time. To probe increases during response preparation, we assessed parametric changes in ERP amplitudes during the interval spanning the final 100 ms prior to response. This interval covered the timeframe of indicated NDT increases, after accounting for the constant probe encoding duration (Figure S3B). Notably, we observed a late frontal potential that increased in amplitude (Figure S3D) and whose onset corresponded to the temporal NDT shift (Figure S3C) after controlling for constant encoding duration (Figure S3B). This suggests that baseline NDT estimates approximate the duration of probe encoding (Nunez, Vandekerckhove, & Srinivasan, 2017), whereas NDT increases characterize increased demands for transforming the sensory decision into a motor command (Lui et al., 2018). This further suggests that drift diffusion modelling successfully dissociated contributions from evidence integration, sensory encoding, and motor preparation. Interestingly, evidence accumulation consistently peaked at/near response execution, suggesting that additional motor demands may unravel in parallel, rather than succeed finished integration (as is often assumed in sequential sampling models).

## Text S4. Behavioral PLS of spectral power during sensation

Task PLS describes the multivariate co-variation of spectral power with load. However, inter-individual behavioral differences may relate to power changes in specific bands. To probe whether inter-individual relations of power modulation to behavior would vary from the mean changes as identified via task PLS, we calculated a behavioral PLS by considering the individual linear change in spectral power with target uncertainty. This revealed a similar multivariate loading pattern as observed for the task PLS (**Figure S4B**), with high agreement between individual brainscores (r = .7, p < .001), suggesting that the identified frequency ranges jointly contributed to behavioral relations.

## Text S5. Pre-stimulus alpha power increases with load, but does not relate to behavioral changes or power changes during sensation

Furthermore, decreases in pre-stimulus alpha power have been linked to increases in cortical excitability at stimulus onset (Iemi, Chaumon, Crouzet, & Busch, 2017; Lange, Oostenveld, & Fries, 2013). To probe whether expected uncertainty modulated pre-stimulus alpha power, we performed another task PLS, covering the final second of the fixation interval prior to stimulus onset. This analysis indicated that pre-stimulus alpha power increased alongside uncertainty (**Figure S4C**). Notably, in contrast to current results, elevated levels of anticipatory alpha power are often associated with decreased gamma power upon stimulus onset. Notably, linear models did not indicate associations between pre-stimulus alpha power increases across load with either drift rate decreases [r(137) = 0.02, 95%CI [−0.15, 0.18], p = 0.86], non-decision time increases [r(137) = 0.06, 95%CI [−0.1, 0.23], p = 0.45] or increases on the SPMF [r(137) = −0.13, 95%CI [−0.29, 0.04], p = 0.13]. These results are in line with increasing evidence suggesting that anticipatory alpha power modulation more closely tracks subjective confidence in upcoming decisions than sensory fidelity (Benwell et al., 2017; Limbach & Corballis, 2016).

## Text S6. SSVEP magnitude is not modulated during sensation

Moreover, SSVEP magnitude has been suggested as a signature of encoded sensory information (O’Connell, Dockree, & Kelly, 2012), that is enhanced by attention (Morgan, Hansen, & Hillyard, 1996; Muller et al., 2006) and indicates fluctuations in excitability (Zhigalov, Herring, Herpers, Bergmann, & Jensen, 2019). However, despite a clear SSVEP signature, we did not observe significant effects of encoding demands on the global SSVEP magnitude (**Figure S4D**). As attentional effects on SSVEP magnitude have been shown to vary by SSVEP frequency (Ding, Sperling, & Srinivasan, 2006), the 30 Hz range may have been suboptimal here. Furthermore, the SSVEP frequency was shared across different features, thus not allowing us to assess whether uncertainty modulated the selective processing of single features. Implementing feature-specific flicker frequencies may overcome such limitations in future work, and allow to assess the changes in feature-specific processing under uncertainty.

## Text S7. Rhythm-specific indices in theta and alpha band relate to multivariate spectral power modulation

Finally, as spectral power conflates rhythmic and arrhythmic signal contributions in magnitude, space and time (Kosciessa, Grandy, Garrett, & Werkle-Bergner, 2020), we performed single-trial rhythm detection, observing similar decreases in the duration and power of alpha rhythms (see **Figure S4E**) that were jointly related to stronger increases on the latent factor [duration: r(137) = −0.61, 95%CI [−0.71, −0.49], p = 1.31e-15; power: r(137) = −0.63, 95%CI [−0.72, −0.52], p = 9.66e-17]. Notably, this analysis indicated increases in theta duration, but not power, suggesting that narrowband theta power changes mainly reflected modulations in the duration of non-stationary theta rhythms, rather than changes in their strength. In line with this suggestion, increases on the spectral power factor related to increases in theta duration [r(137) = 0.19, 95%CI [0.02, 0.35], p = 0.03], but not theta SNR [r(137) = 0.09, 95%CI [−0.08, 0.25], p = 0.31].

## Text S8. A second LV may indicate decreased task engagement due to heightened difficulty at higher uncertainty levels

A 2^nd^ significant LV (p =.012) indicated strong positive loadings in angular gyrus, middle frontal gyrus, and inferior frontal gyrus, as well as occipital cortex (see **Figure S5A**). Negative loadings were observed dominantly in medial PFC, precuneus and V5. This component increased from selective attention to target load 2, but then declined towards higher loads. Decreases in angular gyrus have been strongly to increased visual working memory load (Sheremata, Somers, & Shomstein, 2018; Todd & Marois, 2004). Increases in DMN regions, in addition to decreased prefrontal activity suggest that this component reflects relative task disengagement towards high load conditions, while increases in lateral visual cortex may reflect increased entrainment, and lack of top-down inhibition. In line with more negative loadings on this component being detrimental, we observed that inter-individually higher brainscores (i.e., positive loadings) were associated with lower non-decision times during selective attention (r = −0.46, p = .002), while stronger within-subject decreases with load were associated with larger individual NDT increases [r(122) = −0.18, 95%CI [−0.35, −0.01], p = 0.04] but not changes in drift rate [r(122) = 0.01, 95%CI [−0.17, 0.18], p = 0.95]. Larger decreases on this component were moreover related to more constrained increases in spectral modulation [r(122) = 0.39, 95%CI [0.23, 0.53], p = 6.83e-6]. Jointly, this suggests that individual drop-offs in the positive cluster of regions reflects decreased task engagement under increased difficulty, with adverse behavioral consequences.

---

**Figure S1.**
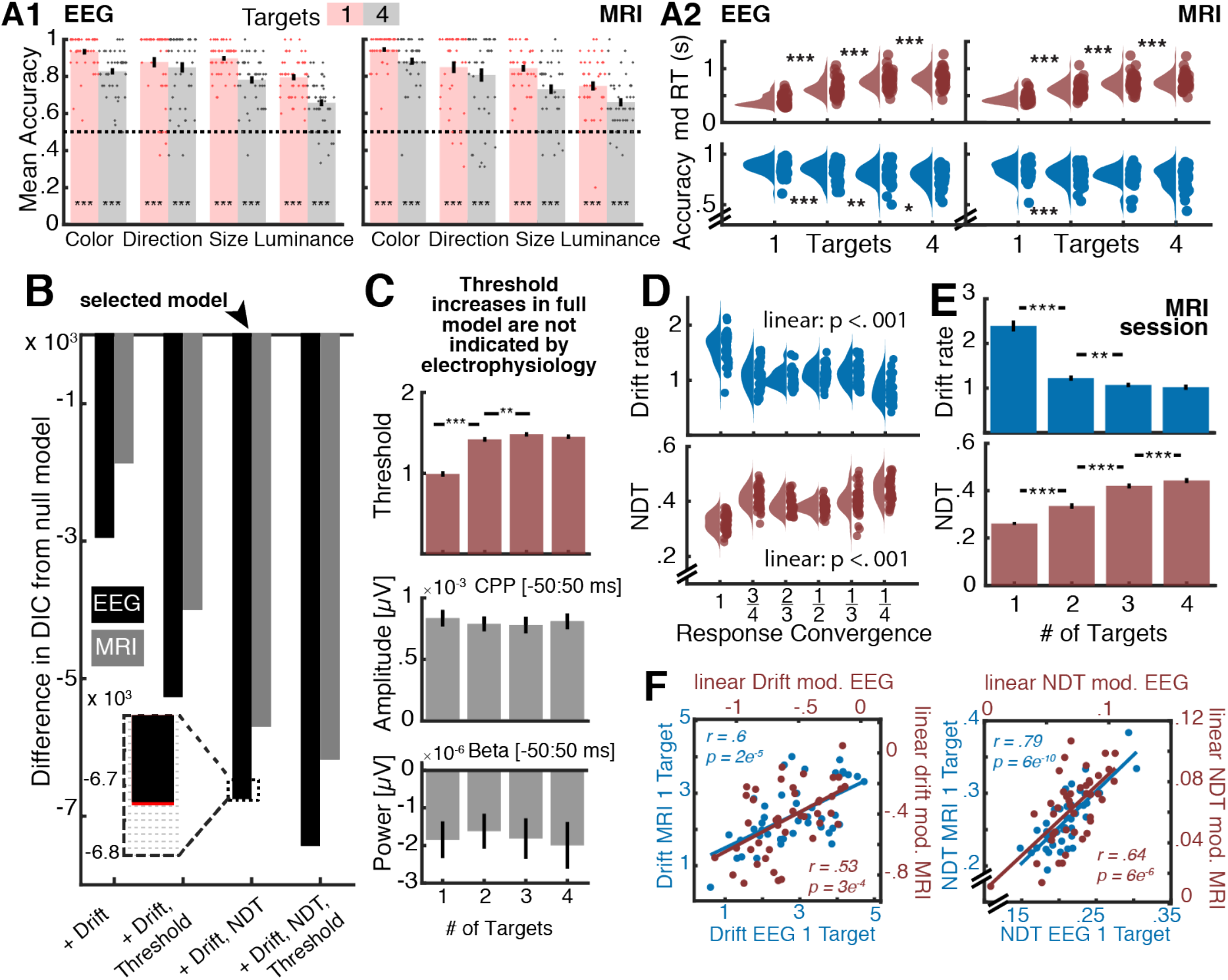
Additional behavioral analyses. **(A1) Accuracies for single target cue and maximum target uncertainty.** For all features, mean accuracy was above chance accuracy (0.5, indicated by broken lines) at the group level. Dots indicate individual accuracies. *** = p < .001 (paired t-test vs. chance accuracy). **(A2) Reaction times and accuracies by load.** All linear effects were significant (p < .001). **(B-C) HDDM model comparison.** (**B**) DIC-based model comparison indicates that full model, including threshold modulation, provides the best group fit to the behavioral data. However, load-related threshold increases (C) were not supported by EEG-based signatures (D). The inset shows an additional comparison of the selected model with an alternative model including starting point variation across load levels (displayed in red). Due to very constrained fit improvements, we selected the simpler model without starting point variation for further analyses. (**C**) **Threshold increases in full model are not indicated by electrophysiology.** The full model indicates additional threshold (also called boundary separation) increases with added target load, with qualitatively identical effects on drift rate and NDT (not shown). Boundary separation captures the conservativeness of the decision criterion and has been related to decision conflict during the choice process (e.g., Cavanagh et al., 2011). EEG-based signatures of evidence integration do not indicate threshold differences. While the full model suggested increased boundary separation, neither of the electrophysiological proxies (i.e., CPP, contralateral beta) of evidence bounds mirrors such increases. While this suggests the absence of threshold increases (McGovern, Hayes, Kelly, & O’Connell, 2018), it alternately questions the sensitivity of electrophysiological threshold estimates, which should be investigated with specific threshold modulations, such as speed-accuracy trade-off instructions, in future work. **(D) Differences in response convergence do not account for main effects of target load.** A separate model including both target load and response convergence indicated practically identical NDT and drift rate effects of target amount, while highlighting additional linear effects of response convergence. Data are individually-centered across conditions. **(E-F) Reliability of individual parameter estimates across sessions.** A separate hierarchical DDM was fit to data from each session. (**E**) Similar group-level effects were indicated for the MRI and EEG (cf. Figure 2B) session: whereas drift rate decreased with load, non-decision time increased. (**F**) Session reliability of inter-individual differences was high both for single-target performance and for linear changes with target load. Reliability was also high for threshold estimates (r = .79, p = 6e-10). [* p <.05; ** p <.01; *** p < .001]

**Figure S2.**
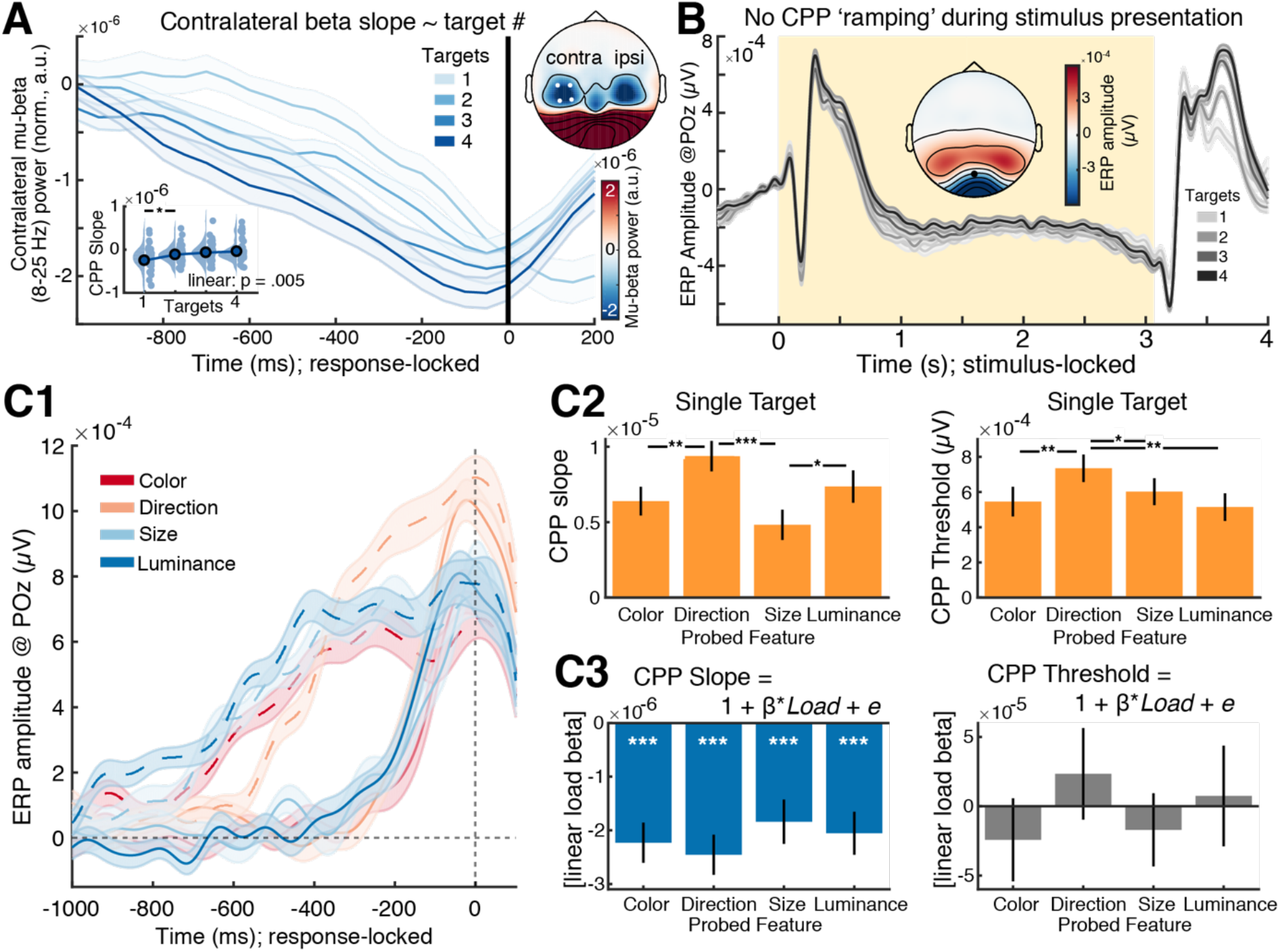
Additional drift rate analyses. **(A) The slope of lateralized motor preparation indicates load-related decreases in drift rate.** (A) Slopes of contralateral mu-beta power shallows with increasing attentional load levels. The inset displays linear slope estimates, estimated via linear regression from −250 ms to −50 ms, relative to response. (B) Topography of response-locked mu-beta power, averaged from −50 ms to +50 ms around response. White dots indicate the contralateral channels from which data was extracted. **(B) The centro-parietal positive potential (CPP) does not show clear ramping increases during stimulus presentation**. The yellow background indicated the stimulus presentation period. Note the modulated ramping following the probe onset at the end of stimulus presentation. The inset shows the topography of the grand average ERPs, temporally averaged during the final 2 seconds of the stimulus presentation period. The black dot indicates channel POz, at which the group-wise CPP was maximal (see Figure 2C1). **(C) Differences between probed stimulus attributes do not account for drift rate decreases under target load.** (A) Response-locked CPP as a function of probed attribute, shown for the single target (complete lines) and four target (broken lines) conditions. Data were selected by condition and probed (cf. cued), attribute, ensuring that unique trials contributed to each load condition. (B) Comparison of CPP slopes and thresholds for different probed features, when the probe target was known in advance. Slopes and thresholds were increased for direction than for other attributes, indicating relatively larger available evidence and more cautious responses (putatively ‘easier’ feature). (C) Load effect of CPP slopes and thresholds for different probed feature attributes. CPP slopes (i.e., evidence drift) exhibited load-related decreases for each probed attribute, whereas no threshold modulation was indicated for any of the probed attributes. [* p <.05; ** p <.01; *** p < .001]

**Figure S3.**
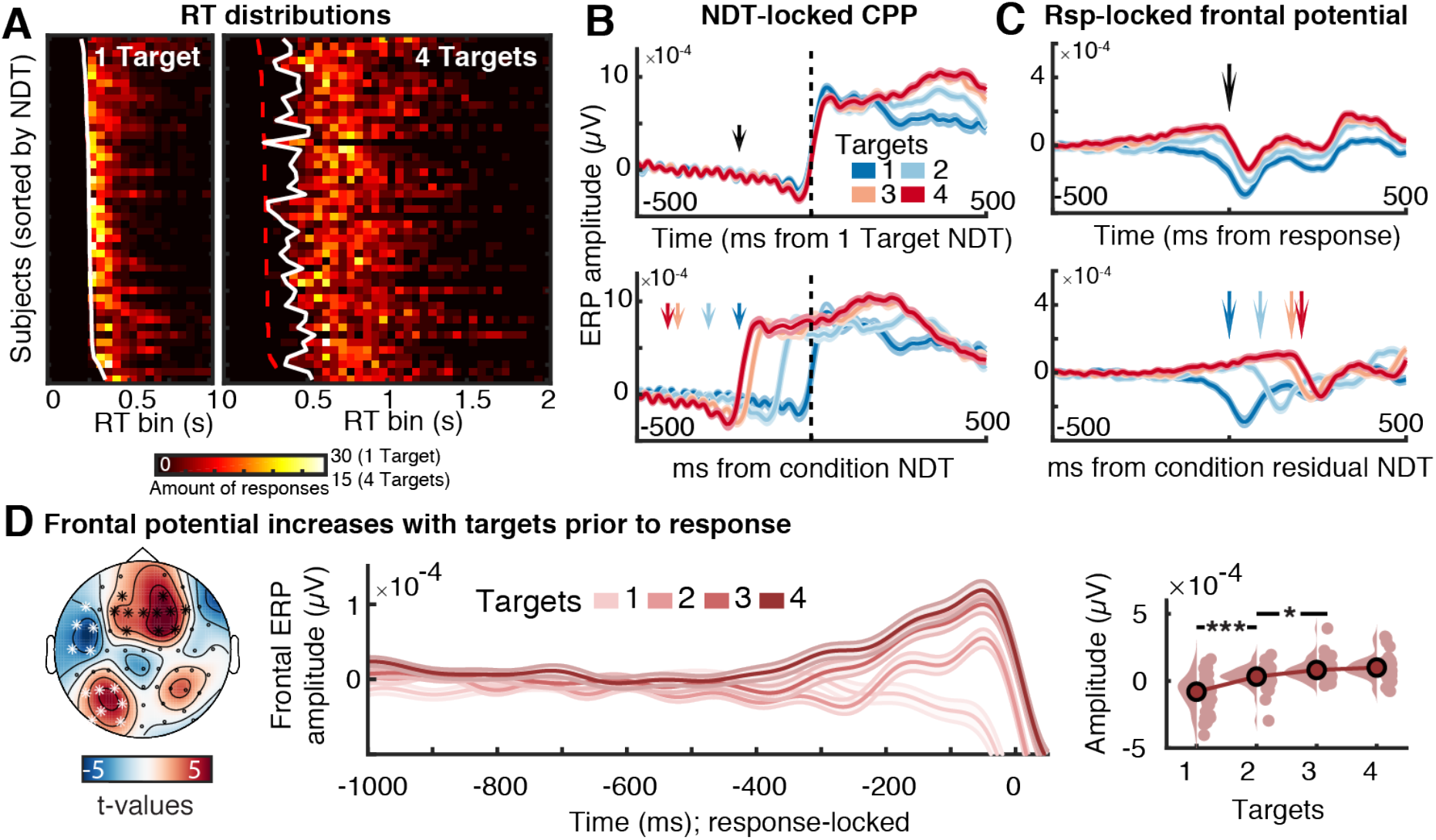
Non-decision time (NDT) increases putatively relate to additional motor demands, not temporal delays in CPP onset. **(A) NDT estimates describe the onset of individual RT distributions (see also Lui et al., 2018).** Response counts (here shown for EEG session) were sorted into 40 bins of 50 ms each. White lines indicate individual NDT estimates; the red dotted line indicates NDT estimates for the single-target condition. **(B, C) Relation of visual encoding and frontal potential to indicated NDT increases.** When response preparation can be made in advance (i.e., when only a single target is indicated) and probe onset only requires response execution, the average NDT estimate aligns with the onset of the CPP (B, top). However, load-related increases in NDT occur in the absence of temporal shifts in CPP onset (B, bottom). In C, arrows indicate the average probe onset time in each condition. In contrast, a frontal potential (see D) increases around the time of residual NDT increases (i.e., NDT estimate for each condition minus constant NDT from single-target condition; C, bottom). In D, arrows indicate the average response time in each condition. **(D) A frontal potential increase prior to response, suggesting that observed NDT increase reflect additional motor preparation demands (e.g., button remapping).** Left: Topography of test for linear ERP changes as a function of load during the final 200 ms prior to response. Clusters in white did not exhibit changes that were exclusive to the period preceding the response (data not shown). Center: Extracted traces averaged within the frontal cluster shown with black asterisks on the left. Right: Post-hoc tests on amplitudes of the frontal potential across the final 100 ms prior to response. Data are individually centered across target loads. [* p <.05; ** p <.01; *** p < .001]

**Figure S4.**
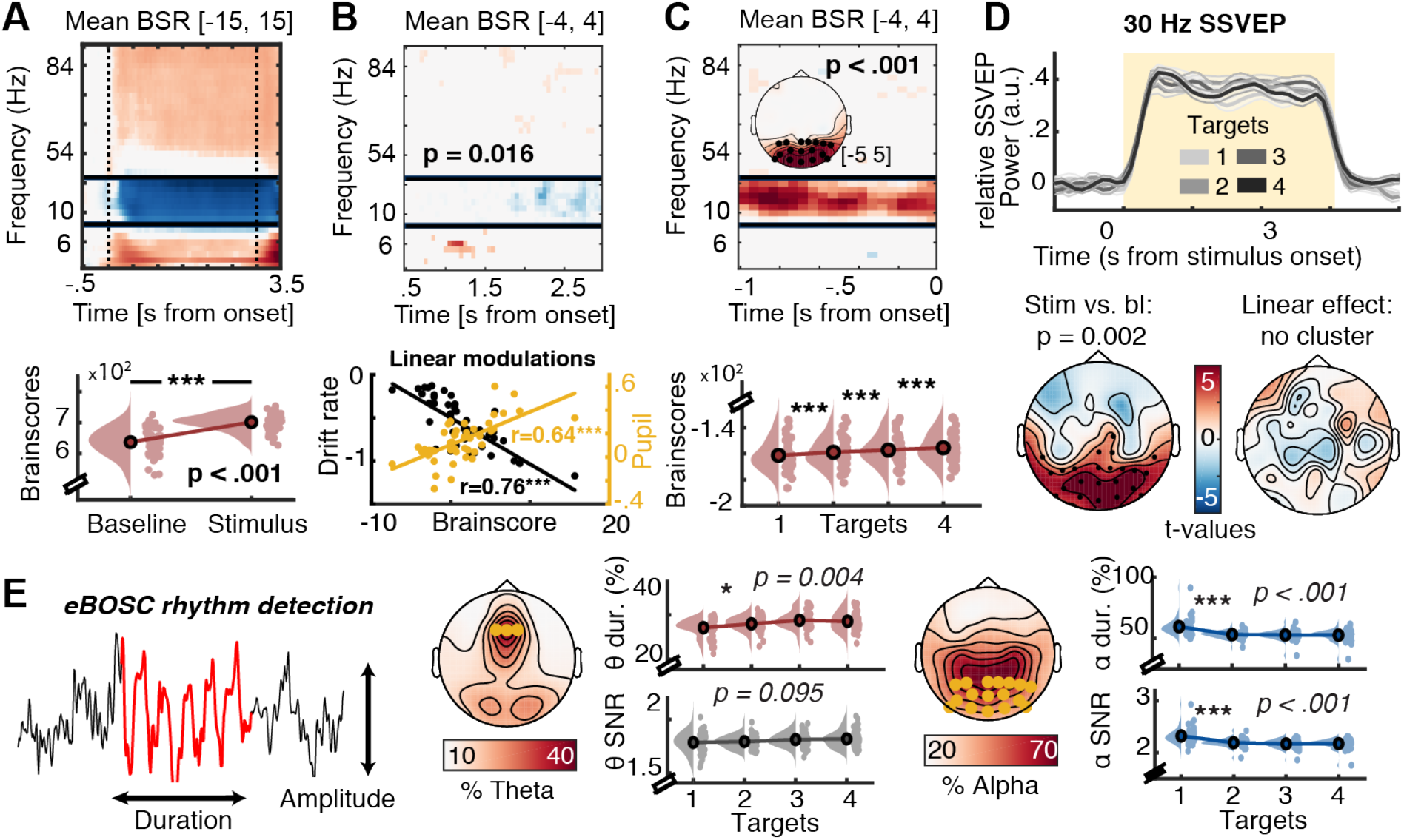
Additional spectral power analyses prior and during sensation. **(A) Multivariate baseline changes and behavioral PLS.** Note that data correspond to the different clusters indicated in Figure 3A. **(B) Behavioral PLS, linking linear multivariate spectral power changes with target # to drift rate decreases and pupil diameter modulation. (C) Parieto-occipital pre-stimulus alpha power increases with target load but is not related to drift changes (see Text S4)**. **(D) SSVEP amplitude is not modulated by attentional load**. Top: Time-resolved, spectrally-normalized, SSVEP power, averaged across occipital channels (O1, Oz, O2), indicates SSVEP presence during stimulus presentation. Bottom left: Topography of stimulus-evoked SSVEP contrast minus baseline. Black dots indicate significant channels as indicated by CBPA. Bottom right: No linear load-related SSVEP modulation was indicated by CBPA. **(E) Modulation of rhythm-specific duration and power by target number.** Left: Schematic of the assessment of amplitude and duration from non-stationary rhythmic events. Right: Topographies of relative theta and alpha occurrence (‘abundance’), averaged across target levels. Orange dots indicate the channels used to extract the data in E, which were the same channels also used in Figure 3AB. Target load decreased alpha duration and power and increased theta duration, but not power. Data are individually centered across target loads. [* p <.05; ** p <.01; *** p < .001]

**Figure S5.**
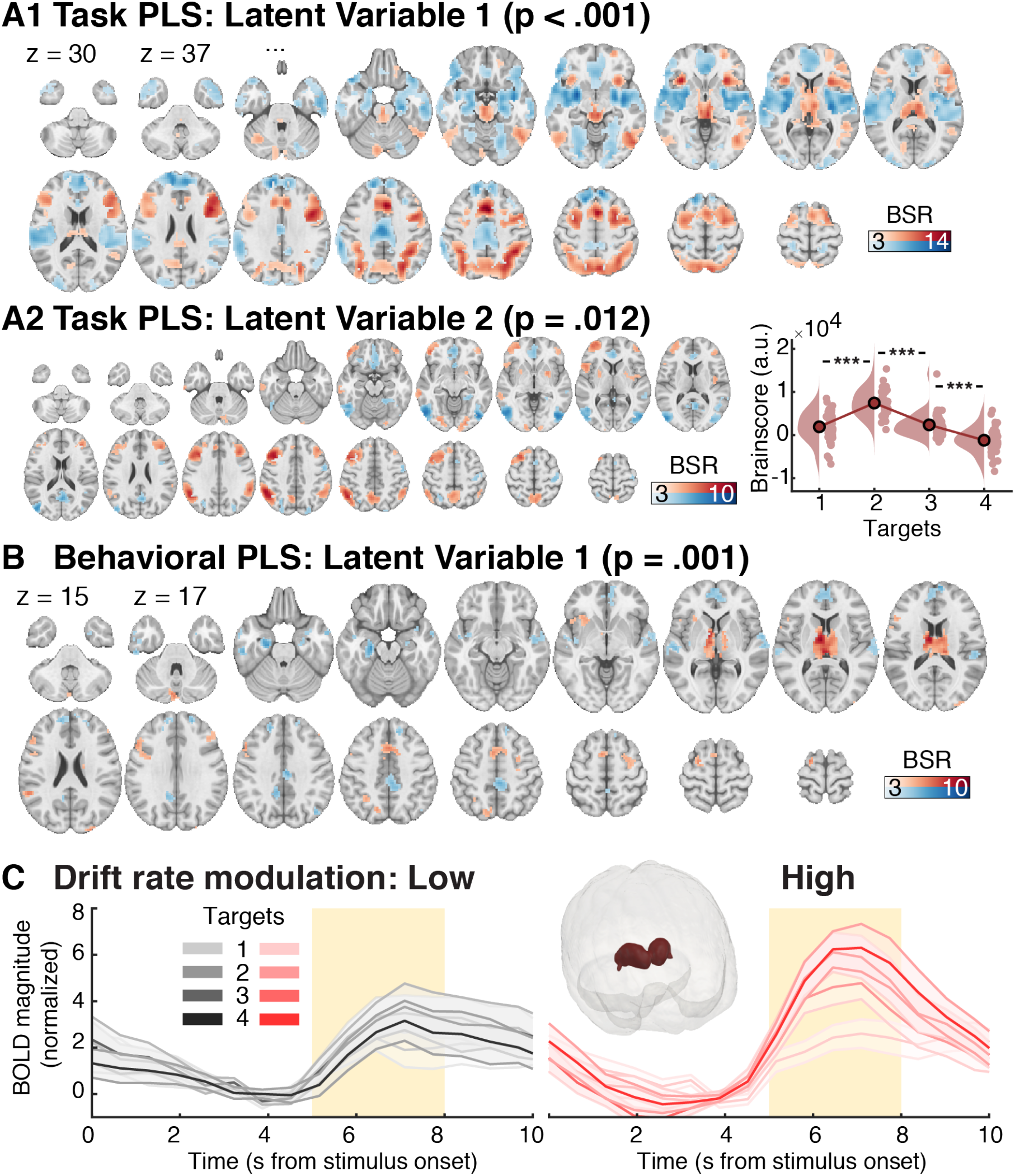
Additional BOLD analyses. **(A, B) Full multivariate brainscore loadings for the two significant latent variables (LVs) produced by the task PLS (A) and behavioral PLS (B).** (A2 left) The brainscore loadings of the second LV designate an initial increase followed by a subsequent decrease towards higher target loads. Data are individually centered across target loads. Thus, the negative components of the pattern expressed on the right become more strongly activated at low and high loads, whereas the positive components are maximally expressed when two targets are relevant. **(C) Thalamic BOLD magnitude for a median split of high- and low drift rate modulators.** The inset shows the thalamic ROI in a glass brain view. [* p <.05; ** p <.01; *** p < .001]

**Table S1.**
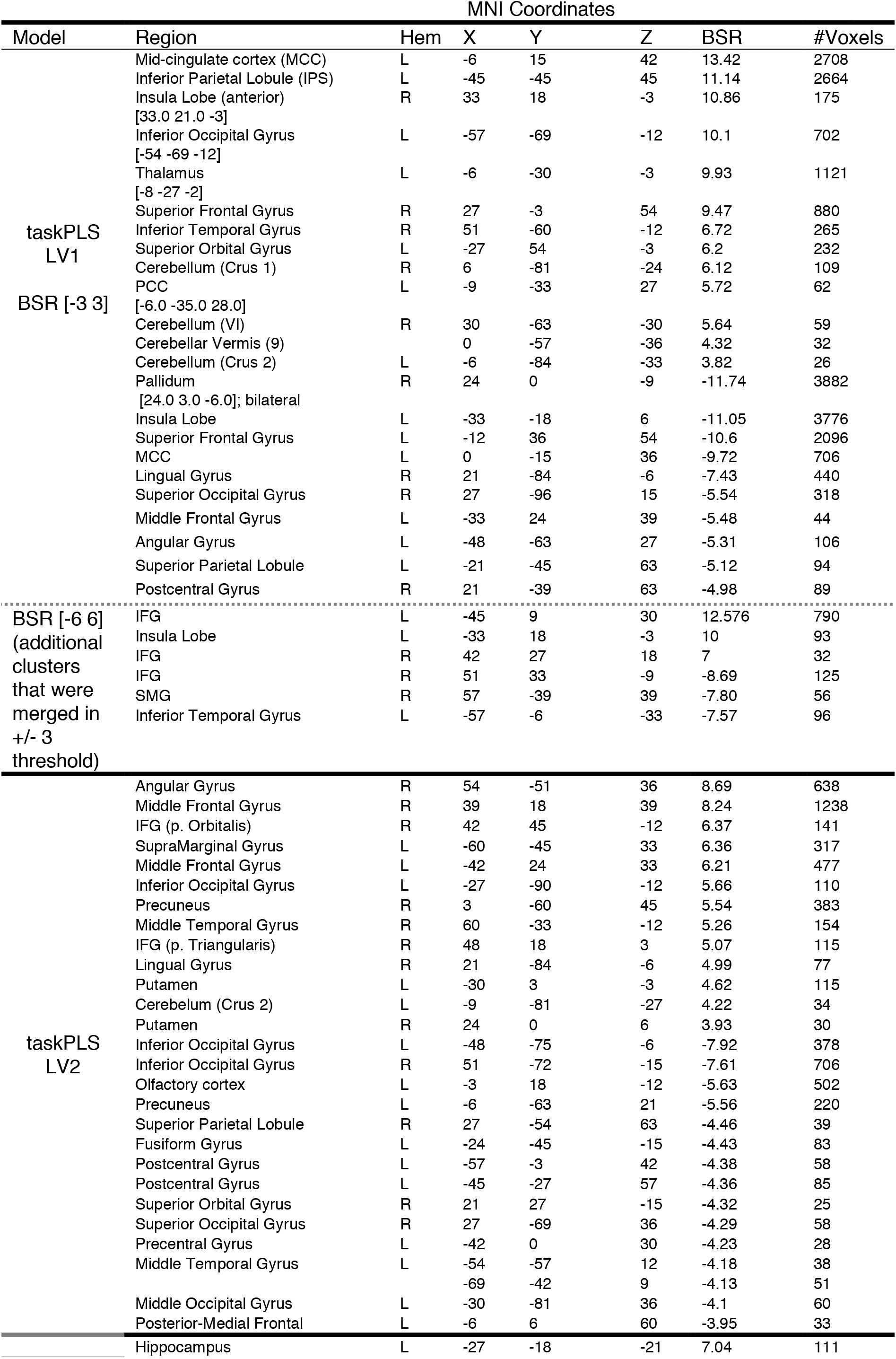
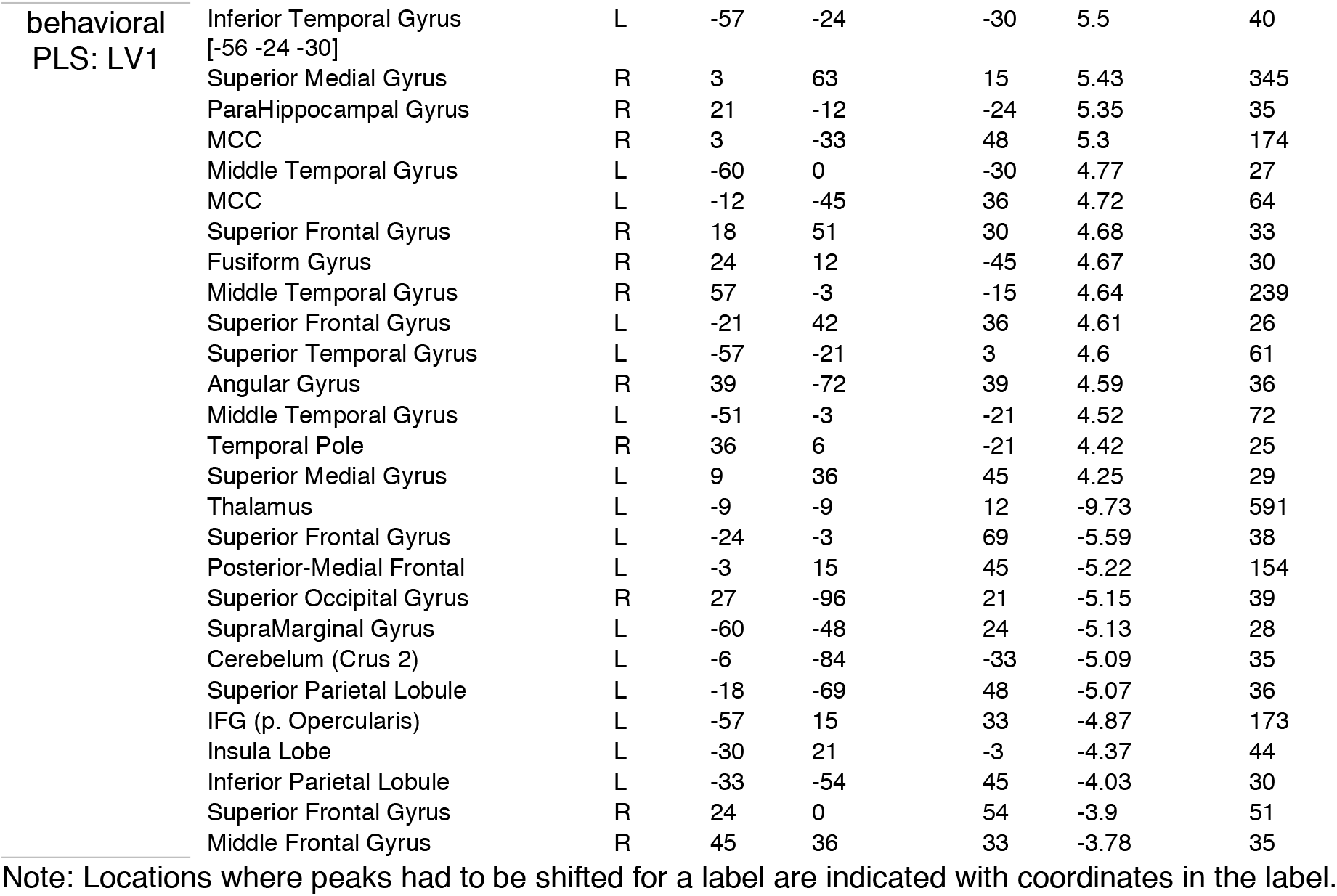
PLS model peak activations, bootstrap ratios, and cluster sizes.

## References

Afyouni, S., & Nichols, T. E. (2018). Insight and inference for DVARS. Neuroimage, 172, 291–312. doi:10.1016/j.neuroimage.2017.12.098

Alitto, H., Rathbun, D. L., Vandeleest, J. J., Alexander, P. C., & Usrey, W. M. (2019). The augmentation of retinogeniculate communication during thalamic burst mode. Journal of Neuroscience, 39(29), 5697–5710. doi:10.1523/Jneurosci.2320-18.2019

Alnaes, D., Sneve, M. H., Espeseth, T., Endestad, T., de Pavert, S. H. P. V., & Laeng, B. (2014). Pupil size signals mental effort deployed during multiple object tracking and predicts brain activity in the dorsal attention network and the locus coeruleus. Journal of Vision, 14(4). doi:10.1167/14.4.1

Arcaro, M. J., Pinsk, M. A., & Kastner, S. (2015). The anatomical and functional organization of the human visual pulvinar. Journal of Neuroscience, 35(27), 9848–9871. doi:10.1523/Jneurosci.1575-14.2015

Aru, J., Aru, J., Priesemann, V., Wibral, M., Lana, L., Pipa, G.,…Vicente, R. (2015). Untangling cross-frequency coupling in neuroscience. Current Opinion in Neurobiology, 31, 51–61. doi:10.1016/j.conb.2014.08.002

Aston-Jones, G., & Cohen, J. D. (2005). An integrative theory of locus coeruleus-norepinephrine function: Adaptive gain and optimal performance. Annual Review of Neuroscience, 28, 403–450. doi:10.1146/annurev.neuro.28.061604.135709

Atallah, B. V., & Scanziani, M. (2009). Instantaneous modulation of gamma oscillation frequency by balancing excitation with inhibition. Neuron, 62(4), 566–577. doi:10.1016/j.neuron.2009.04.027

Avants, B. B., Tustison, N. J., Song, G., Cook, P. A., Klein, A., & Gee, J. C. (2011). A reproducible evaluation of ANTs similarity metric performance in brain image registration. Neuroimage, 54(3), 2033–2044. doi:10.1016/j.neuroimage.2010.09.025

Bach, D. R., & Dolan, R. J. (2012). Knowing how much you don’t know: A neural organization of uncertainty estimates. Nature Reviews Neuroscience, 13(8), 572–586. doi:10.1038/nrn3289

Bakdash, J. Z., & Marusich, L. R. (2017). Repeated measures correlation. Frontiers in Psychology, 8. doi:10.3389/fpsyg.2017.00456

Banca, P., Vestergaard, M. D., Rankov, V., Baek, K., Mitchell, S., Lapa, T.,…Voon, V. (2015). Evidence accumulation in obsessive-compulsive disorder: The role of uncertainty and monetary reward on perceptual decision-making thresholds. Neuropsychopharmacology, 40(5), 1192–1202. doi:10.1038/npp.2014.303

Bauer, M., Kluge, C., Bach, D., Bradbury, D., Heinze, H. J., Dolan, R. J., & Driver, J. (2012). Cholinergic enhancement of visual attention and neural oscillations in the human brain. Current Biology, 22(5), 397–402. doi:10.1016/j.cub.2012.01.022

Beckmann, C. F., & Smith, S. A. (2004). Probabilistic independent component analysis for functional magnetic resonance imaging. Ieee Transactions on Medical Imaging, 23(2), 137–152. doi:10.1109/Tmi.2003.822821

Bell, A. J., & Sejnowski, T. J. (1995). An information maximization approach to blind separation and blind deconvolution. Neural Computation, 7(6), 1129–1159. doi:10.1162/neco.1995.7.6.1129

Berridge, C. W., & Waterhouse, B. D. (2003). The locus coeruleus-noradrenergic system: Modulation of behavioral state and state-dependent cognitive processes. Brain Research Reviews, 42(1), 33–84. doi:10.1016/S0165-0173(03)00143-7

Billig, A. J., Herrmann, B., Rhone, A. E., Gander, P. E., Nourski, K. V., Snoad, B. F.,…Johnsrude, I. S. (2019). A sound-sensitive source of alpha oscillations in human non-primary auditory cortex. Journal of Neuroscience, 39(44), 8679–8689. doi:10.1523/Jneurosci.0696-19.2019

Birn, R. M. (2012). The role of physiological noise in resting-state functional connectivity. Neuroimage, 62(2), 864–870. doi:10.1016/j.neuroimage.2012.01.016

Brainard, D. H. (1997). The psychophysics toolbox. Spatial Vision, 10(4), 433–436. doi:10.1163/156856897x00357

Breton-Provencher, V., & Sur, M. (2019). Active control of arousal by a locus coeruleus GABAergic circuit. Nature Neuroscience, 22(2), 218–228. doi:10.1038/s41593-018-0305-z

Buschman, T. J., & Kastner, S. (2015). From behavior to neural dynamics: An integrated theory of attention. Neuron, 88(1), 127–144. doi:10.1016/j.neuron.2015.09.017

Canolty, R. T., Edwards, E., Dalal, S. S., Soltani, M., Nagarajan, S. S., Kirsch, H. E.,…Knight, R. T. (2006). High gamma power is phase-locked to theta oscillations in human neocortex. Science, 313(5793), 1626–1628. doi:10.1126/science.1128115

Caplan, J. B., Madsen, J. R., Raghavachari, S., & Kahana, M. J. (2001). Distinct patterns of brain oscillations underlie two basic parameters of human maze learning. Journal of Neurophysiology, 86(1), 368–380.

Cavanagh, J. F., & Frank, M. J. (2014). Frontal theta as a mechanism for cognitive control. Trends in Cognitive Sciences, 18(8), 414–421. doi:10.1016/j.tics.2014.04.012

Cheung, M. J., Kovacevic, N., Fatima, Z., Misic, B., & McIntosh, A. R. (2016). [MEG]PLS: A pipeline for MEG data analysis and partial least squares statistics. Neuroimage, 124, 181–193. doi:10.1016/j.neuroimage.2015.08.045

Colombo, M. A., Napolitani, M., Boly, M., Gosseries, O., Casarotto, S., Rosanova, M.,…Sarasso, S. (2019). The spectral exponent of the resting EEG indexes the presence of consciousness during unresponsiveness induced by propofol, xenon, and ketamine. Neuroimage, 189, 631–644. doi:10.1016/j.neuroimage.2019.01.024

Constantinople, C. M., & Bruno, R. M. (2011). Effects and mechanisms of wakefulness on local cortical networks. Neuron, 69(6), 1061–1068. doi:10.1016/j.neuron.2011.02.040

Crick, F. (2003). Function of the thalamic reticular complex: The searchlight hypothesis. Essential Sources in the Scientific Study of Consciousness, 263–272.

Dahl, M. J., Mather, M., Sander, M. C., & Werkle-Bergne, M. (2020). Noradrenergic responsiveness supports selective attention across the adult lifespan. Journal of Neuroscience, 40(22), 4372–4390. doi:10.1523/Jneurosci.0398-19.2020

de Gee, J. W., Colizoli, O., Kloosterman, N. A., Knapen, T., Nieuwenhuis, S., & Donner, T. H. (2017). Dynamic modulation of decision biases by brainstem arousal systems. Elife, 6. doi:10.7554/eLife.23232

Dehghani, N., & Wimmer, R. D. (2019). A computational perspective of the role of the thalamus in cognition. Neural Computation, 31(7), 1380–1418. doi:10.1162/neco_a_01197

Desimone, R., & Duncan, J. (1995). Neural mechanisms of selective visual-attention. Annual Review of Neuroscience, 18, 193–222. doi:10.1146/annurev.ne.18.030195.001205

Destexhe, A., & Rudolph, M. (2004). Extracting information from the power spectrum of synaptic noise. Journal of Computational Neuroscience, 17(3), 327–345. doi:Doi 10.1023/B:Jcns.0000044875.90630.88

Destexhe, A., Rudolph, M., & Pare, D. (2003). The high-conductance state of neocortical neurons in vivo. Nature Reviews Neuroscience, 4(9), 739–751. doi:10.1038/nrn1198

Ding, J., Sperling, G., & Srinivasan, R. (2006). Attentional modulation of SSVEP power depends on the network tagged by the flicker frequency. Cerebral Cortex, 16(7), 1016–1029. doi:10.1093/cercor/bhj044

Donner, T. H., Siegel, M., Fries, P., & Engel, A. K. (2009). Buildup of choice-predictive activity in human motor cortex during perceptual decision making. Current Biology, 19(18), 1581–1585. doi:10.1016/j.cub.2009.07.066

Dosenbach, N. U., Fair, D. A., Miezin, F. M., Cohen, A. L., Wenger, K. K., Dosenbach, R. A.,…Petersen, S. E. (2007). Distinct brain networks for adaptive and stable task control in humans. Proc Natl Acad Sci U S A, 104(26), 11073–11078. doi:10.1073/pnas.0704320104

Dube, B., Emrich, S. M., & Al-Aidroos, N. (2017). More than a filter: Feature-based attention regulates the distribution of visual working memory resources. Journal of Experimental Psychology-Human Perception and Performance, 43(10), 1843–1854. doi:10.1037/xhp0000428

Dugue, L., Marque, P., & VanRullen, R. (2011). The phase of ongoing oscillations mediates the causal relation between brain excitation and visual perception. Journal of Neuroscience, 31(33), 11889–11893. doi:10.1523/Jneurosci.1161-11.2011

Efron, B., & Tibshirani, R. (1986). Bootstrap methods for standard errors, confidence intervals, and other measures of statistical accuracy. Statist. Sci., 1(1), 54–75. doi:10.1214/ss/1177013815

Eickhoff, S. B., Stephan, K. E., Mohlberg, H., Grefkes, C., Fink, G. R., Amunts, K., & Zilles, K. (2005). A new SPM toolbox for combining probabilistic cytoarchitectonic maps and functional imaging data. Neuroimage, 25(4), 1325–1335. doi:10.1016/j.neuroimage.2004.12.034

Ferguson, K. A., & Cardin, J. A. (2020). Mechanisms underlying gain modulation in the cortex. Nature Reviews Neuroscience, 21(2), 80–92. doi:10.1038/s41583-019-0253-y

Fiebelkorn, I. C., Pinsk, M. A., & Kastner, S. (2019). The mediodorsal pulvinar coordinates the macaque fronto-parietal network during rhythmic spatial attention. Nature Communications, 10. doi:10.1038/s41467-018-08151-4

Fonov, V., Evans, A. C., Botteron, K., Almli, C. R., McKinstry, R. C., Collins, D. L., & Grp, B. C. (2011). Unbiased average age-appropriate atlases for pediatric studies. Neuroimage, 54(1), 313–327. doi:10.1016/j.neuroimage.2010.07.033

Forstmann, B. U., Ratcliff, R., & Wagenmakers, E. J. (2016). Sequential sampling models in cognitive neuroscience: Advantages, applications, and extensions. Annual Review of Psychology, Vol 67, 67, 641–666. doi:10.1146/annurev-psych-122414-033645

Frank, M. J., Gagne, C., Nyhus, E., Masters, S., Wiecki, T. V., Cavanagh, J. F., & Badre, D. (2015). fMRI and EEG predictors of dynamic decision parameters during human reinforcement learning. Journal of Neuroscience, 35(2), 485–494. doi:10.1523/Jneurosci.2036-14.2015

Fries, P. (2015). Rhythms for cognition: Communication through coherence. Neuron, 88(1), 220–235. doi:10.1016/j.neuron.2015.09.034

Friston, K. J., Williams, S., Howard, R., Frackowiak, R. S., & Turner, R. (1996). Movement-related effects in fMRI time-series. Magn Reson Med, 35(3), 346–355. doi:10.1002/mrm.1910350312

Froemke, R. C. (2015). Plasticity of cortical excitatory-inhibitory balance. Annual Review of Neuroscience, 38, 195–219. doi:10.1146/annurev-neuro-071714-034002

Froemke, R. C., Merzenich, M. M., & Schreiner, C. E. (2007). A synaptic memory trace for cortical receptive field plasticity. Nature, 450(7168), 425–429. doi:10.1038/nature06289

Furey, M. L., Pietrini, P., & Haxby, J. V. (2000). Cholinergic enhancement and increased selectivity of perceptual processing during working memory. Science, 290(5500), 2315–2319. doi:10.1126/science.290.5500.2315

Gao, R., Peterson, E. J., & Voytek, B. (2017). Inferring synaptic excitation/inhibition balance from field potentials. Neuroimage, 158, 70–78. doi:10.1016/j.neuroimage.2017.06.078

Garrett, D. D., Epp, S. M., Perry, A., & Lindenberger, U. (2018). Local temporal variability reflects functional integration in the human brain. Neuroimage, 183, 776–787. doi:10.1016/j.neuroimage.2018.08.019

Garrett, D. D., Kovacevic, N., McIntosh, A. R., & Grady, C. L. (2010). Blood oxygen level-dependent signal variability is more than just noise. Journal of Neuroscience, 30(14), 4914–4921. doi:10.1523/Jneurosci.5166-09.2010

Garrett, D. D., McIntosh, A. R., & Grady, C. L. (2014). Brain signal variability is parametrically modifiable. Cerebral Cortex, 24(11), 2931–2940. doi:10.1093/cercor/bht150

Garrett, D. D., Nagel, I. E., Preuschhof, C., Burzynska, A. Z., Marchner, J., Wiegert, S.,…Lindenberger, U. (2015). Amphetamine modulates brain signal variability and working memory in younger and older adults. Proceedings of the National Academy of Sciences of the United States of America, 112(24), 7593–7598. doi:10.1073/pnas.1504090112

Gold, J. I., & Shadlen, M. N. (2007). The neural basis of decision making. Annual Review of Neuroscience, 30, 535–574. doi:10.1146/annurev.neuro.29.051605.113038

Grandy, T. H., Garrett, D. D., Schmiedek, F., & Werkle-Bergner, M. (2016). On the estimation of brain signal entropy from sparse neuroimaging data. Scientific Reports, 6. doi:10.1038/srep23073

Haegens, S., Nacher, V., Luna, R., Romo, R., & Jensen, O. (2011). alpha-Oscillations in the monkey sensorimotor network influence discrimination performance by rhythmical inhibition of neuronal spiking. Proceedings of the National Academy of Sciences of the United States of America, 108(48), 19377–19382. doi:10.1073/pnas.1117190108

Halassa, M. M., & Kastner, S. (2017). Thalamic functions in distributed cognitive control. Nature Neuroscience, 20(12), 1669–1679. doi:10.1038/s41593-017-0020-1

Halassa, M. M., & Sherman, S. M. (2019). Thalamocortical circuit motifs: A general framework. Neuron, 103(5), 762–770. doi:10.1016/j.neuron.2019.06.005

Halgren, M., Ulbert, I., Bastuji, H., Fabo, D., Eross, L., Rey, M.,…Cash, S. S. (2019). The generation and propagation of the human alpha rhythm. Proceedings of the National Academy of Sciences of the United States of America, 116(47), 23772–23782. doi:10.1073/pnas.1913092116

Hampshire, A., Chamberlain, S. R., Monti, M. M., Duncan, J., & Owen, A. M. (2010). The role of the right inferior frontal gyrus: Inhibition and attentional control. Neuroimage, 50(3), 1313–1319. doi:10.1016/j.neuroimage.2009.12.109

Hanks, T. D., & Summerfield, C. (2017). Perceptual decision making in rodents, monkeys, and humans. Neuron, 93(1), 15–31. doi:10.1016/j.neuron.2016.12.003

Harris, K. D., & Thiele, A. (2011). Cortical state and attention. Nature Reviews Neuroscience, 12(9), 509–523. doi:10.1038/nrn3084

Hartings, J. A., Temereanca, S., & Simons, D. J. (2003). State-dependent processing of sensory stimuli by thalamic reticular neurons. Journal of Neuroscience, 23(12), 5264–5271.

Hirata, A., Aguilar, J., & Castro-Alamancos, M. A. (2006). Noradrenergic activation amplifies bottom-up and top-down signal-to-noise ratios in sensory thalamus. Journal of Neuroscience, 26(16), 4426–4436. doi:10.1523/JNEUROSCI.5298-05.2006

Honjoh, S., Sasai, S., Schiereck, S. S., Nagai, H., Tononi, G., & Cirelli, C. (2018). Regulation of cortical activity and arousal by the matrix cells of the ventromedial thalamic nucleus. Nature Communications, 9. doi:10.1038/s41467-018-04497-x

Horn, A., & Blankenburg, F. (2016). Toward a standardized structural-functional group connectome in mni space. Neuroimage, 124, 310–322. doi:10.1016/j.neuroimage.2015.08.048

Hwang, K., Bertolero, M. A., Liu, W. B., & D’Esposito, M. (2017). The human thalamus is an integrative hub for functional brain networks. Journal of Neuroscience, 37(23), 5594–5607. doi:10.1523/Jneurosci.0067-17.2017

Iemi, L., Chaumon, M., Crouzet, S. M., & Busch, N. A. (2017). Spontaneous neural oscillations bias perception by modulating baseline excitability. Journal of Neuroscience, 37(4), 807–819. doi:10.1523/JNEUROSCI.1432-16.2016

Jagtap, P., & Diwadkar, V. A. (2016). Effective connectivity of ascending and descending frontalthalamic pathways during sustained attention: Complex brain network interactions in adolescence. Human Brain Mapping, 37(7), 2557–2570. doi:10.1002/hbm.23196

Jaramillo, J., Mejias, J. F., & Wang, X. J. (2019). Engagement of pulvino-cortical feedforward and feedback pathways in cognitive computations. Neuron, 101(2), 321–336. doi:10.1016/j.neuron.2018.11.023

Jasper, H. H. (1948). Charting the sea of brain waves. Science, 108(2805), 343–347. doi:10.1126/science.108.2805.343

Jenkinson, M., Beckmann, C. F., Behrens, T. E., Woolrich, M. W., & Smith, S. M. (2012). Fsl. Neuroimage, 62(2), 782–790. doi:10.1016/j.neuroimage.2011.09.015

Jones, E. G. (2009). Synchrony in the interconnected circuitry of the thalamus and cerebral cortex. Disorders of Consciousness, 1157, 10–23. doi:10.1111/j.1749-6632.2009.04534.x

Joshi, S., & Gold, J. I. (2020). Pupil size as a window on neural substrates of cognition. Trends in Cognitive Sciences, 24(6), 466–480. doi:10.1016/j.tics.2020.03.005

Joshi, S., Li, Y., Kalwani, R. M., & Gold, J. I. (2016). Relationships between pupil diameter and neuronal activity in the locus coeruleus, colliculi, and cingulate cortex. Neuron, 89(1), 221–234. doi:10.1016/j.neuron.2015.11.028

Kanai, R., Komura, Y., Shipp, S., & Friston, K. (2015). Cerebral hierarchies: Predictive processing, precision and the pulvinar. Philosophical Transactions of the Royal Society B-Biological Sciences, 370(1668), 69–81. doi:10.1098/rstb.2014.0169

Kelly, S. P., & O’Connell, R. G. (2013). Internal and external influences on the rate of sensory evidence accumulation in the human brain. Journal of Neuroscience, 33(50), 19434–19441. doi:10.1523/Jneurosci.3355-13.2013

Kim, C., Cilles, S. E., Johnson, N. F., & Gold, B. T. (2012). Domain general and domain preferential brain regions associated with different types of task switching: A meta-analysis. Human Brain Mapping, 33(1), 130–142. doi:10.1002/hbm.21199

Kleiner, M., Brainard, D., & Pelli, D. (2007). What’s new in psychtoolbox-3? Perception, 36, 14–14.

Klimesch, W., Sauseng, P., & Hanslmayr, S. (2007). EEG alpha oscillations: The inhibition-timing hypothesis. Brain Res Rev, 53(1), 63–88. doi:10.1016/j.brainresrev.2006.06.003

Kloosterman, N. A., Kosciessa, J. Q., Lindenberger, U., Fahrenfort, J. J., & Garrett, D. D. (2019). Boosting brain signal variability underlies liberal shifts in decision bias. bioRxiv.

Koelewijn, T., Shinn-Cunningham, B. G., Zekveld, A. A., & Kramer, S. E. (2014). The pupil response is sensitive to divided attention during speech processing. Hearing Research, 312, 114–120. doi:10.1016/j.heares.2014.03.010

Kosciessa, J. Q., Grandy, T. H., Garrett, D. D., & Werkle-Bergner, M. (2020). Single-trial characterization of neural rhythms: Potential and challenges. Neuroimage, 206, 116331. doi:10.1016/j.neuroimage.2019.116331

Kosciessa, J. Q., Kloosterman, N. A., & Garrett, D. D. (2020). Standard multiscale entropy reflects neural dynamics at mismatched temporal scales: What’s signal irregularity got to do with it? Plos Computational Biology, 16(5), e1007885. doi:10.1371/journal.pcbi.1007885

Krishnamurthy, K., Nassar, M. R., Sarode, S., & Gold, J. I. (2017). Arousal-related adjustments of perceptual biases optimize perception in dynamic environments. Nature Human Behaviour, 1(6). doi:10.1038/s41562-017-0107

Krishnan, A., Williams, L. J., McIntosh, A. R., & Abdi, H. (2011). Partial least squares (PLS) methods for neuroimaging: A tutorial and review. Neuroimage, 56(2), 455–475. doi:10.1016/j.neuroimage.2010.07.034

Lange, J., Oostenveld, R., & Fries, P. (2013). Reduced occipital alpha power indexes enhanced excitability rather than improved visual perception. Journal of Neuroscience, 33(7), 3212–3220. doi:10.1523/Jneurosci.3755-12.2013

Lee, S. H., & Dan, Y. (2012). Neuromodulation of brain states. Neuron, 76(1), 209–222. doi:10.1016/j.neuron.2012.09.012

Lendner, J. D., Helfrich, R. F., Mander, B. A., Romundstad, L., Lin, J. J., Walker, M. P.,…Knight, R. T. (2019). An electrophysiological marker of arousal level in humans. bioRxiv.

Lewis, L. D., Voigts, J., Flores, F. J., Schmitt, L. I., Wilson, M. A., Halassa, M. M., & Brown, N. (2015). Thalamic reticular nucleus induces fast and local modulation of arousal state. Elife, 4, e08760. doi:10.7554/eLife.08760

Liu, J., Lee, H. J., Weitz, A. J., Fang, Z. N., Lin, P., Choy, M.,…Lee, J. H. (2015). Frequency-selective control of cortical and subcortical networks by central thalamus. Elife, 4. doi:10.7554/eLife.09215

Lopes da Silva, F. H., Vos, J. E., Mooibroek, J., & Van Rotterdam, A. (1980). Relative contributions of intracortical and thalamo-cortical processes in the generation of alpha rhythms, revealed by partial coherence analysis. Electroencephalogr Clin Neurophysiol, 50(5-6), 449–456. doi:10.1016/0013-4694(80)90011-5

Lorincz, M. L., Kekesi, K. A., Juhasz, G., Crunelli, V., & Hughes, S. W. (2009). Temporal framing of thalamic relay-mode firing by phasic inhibition during the alpha rhythm. Neuron, 63(5), 683–696. doi:10.1016/j.neuron.2009.08.012

Marguet, S. L., & Harris, K. D. (2011). State-dependent representation of amplitude-modulated noise stimuli in rat auditory cortex. Journal of Neuroscience, 31(17), 6414–6420. doi:10.1523/Jneurosci.5773-10.2011

Maris, E., & Oostenveld, R. (2007). Nonparametric statistical testing of EEG- and MEG-data. Journal of Neuroscience Methods, 164(1), 177–190. doi:10.1016/j.jneumeth.2007.03.024

Martins, A. R. O., & Froemke, R. C. (2015). Coordinated forms of noradrenergic plasticity in the locus coeruleus and primary auditory cortex. Nature Neuroscience, 18(10), 1483–1492. doi:10.1038/nn.4090

Marton, T. F., Seifikar, H., Luongo, F. J., Lee, A. T., & Sohal, V. S. (2018). Roles of prefrontal cortex and mediodorsal thalamus in task engagement and behavioral flexibility. Journal of Neuroscience, 38(10), 2569–2578. doi:10.1523/Jneurosci.1728-17.2018

Maunsell, J. H. R. (2015). Neuronal mechanisms of visual attention. Annual Review of Vision Science, Vol 1, 1, 373–391. doi:10.1146/annurev-vision-082114-035431

McCormick, D. A. (1989). Cholinergic and noradrenergic modulation of thalamocortical processing. Trends in Neurosciences, 12(6), 215–221. doi:10.1016/0166-2236(89)90125-2

McCormick, D. A., McGinley, M. J., & Salkoff, D. B. (2015). Brain state dependent activity in the cortex and thalamus. Current Opinion in Neurobiology, 31, 133–140. doi:10.1016/j.conb.2014.10.003

McCormick, D. A., Pape, H. C., & Williamson, A. (1991). Actions of norepinephrine in the cerebral cortex and thalamus: Implications for function of the central noradrenergic system. Prog Brain Res, 88, 293–305. doi:10.1016/s0079-6123(08)63817-0

McFadyen, J., Dolan, R. J., & Garrido, M. I. (2020). The influence of subcortical shortcuts on disordered sensory and cognitive processing. Nature Reviews Neuroscience, 21(5), 264–276. doi:10.1038/s41583-020-0287-1

McGinley, M. J., David, S. V., & McCormick, D. A. (2015). Cortical membrane potential signature of optimal states for sensory signal detection. Neuron, 87(1), 179–192. doi:10.1016/j.neuron.2015.05.038

McGinley, M. J., Vinck, M., Reimer, J., Batista-Brito, R., Zagha, E., Cadwell, C. R.,…McCormick, D. A. (2015). Waking state: Rapid variations modulate neural and behavioral responses. Neuron, 87(6), 1143–1161. doi:10.1016/j.neuron.2015.09.012

McGovern, D. P., Hayes, A., Kelly, S. P., & O’Connell, R. G. (2018). Reconciling age-related changes in behavioural and neural indices of human perceptual decision-making. Nature Human Behaviour, 2(12), 955–966. doi:10.1038/s41562-018-0465-6

McIntosh, A. R., Bookstein, F. L., Haxby, J. V., & Grady, C. L. (1996). Spatial pattern analysis of functional brain images using partial least squares. Neuroimage, 3(3), 143–157. doi:10.1006/nimg.1996.0016

McIntosh, A. R., & Lobaugh, N. J. (2004). Partial least squares analysis of neuroimaging data: Applications and advances. Neuroimage, 23, S250–S263. doi:10.1016/j.neuroimage.2004.07.020

Minces, V., Pinto, L., Dan, Y., & Chiba, A. A. (2017). Cholinergic shaping of neural correlations. Proceedings of the National Academy of Sciences of the United States of America, 114(22), 5725–5730. doi:10.1073/pnas.1621493114

Mo, C., Lu, J. S., Wu, B. C., Jia, J. R., Luo, H., & Fang, F. (2019). Competing rhythmic neural representations of orientations during concurrent attention to multiple orientation features. Nature Communications, 10. doi:10.1038/s41467-019-13282-3

Murphy, P. R., Wilming, N., Hernandez-Bocanegra, D. C., Prat Ortega, G., & Donner, T. H. (2020). Normative circuit dynamics across human cortex during evidence accumulation in changing environments. bioRxiv.

Nassar, M. R., Rumsey, K. M., Wilson, R. C., Parikh, K., Heasly, B., & Gold, J. I. (2012). Rational regulation of learning dynamics by pupil-linked arousal systems. Nature Neuroscience, 15(7), 1040–1046. doi:10.1038/nn.3130

Nelson, S. M., Dosenbach, N. U. F., Cohen, A. L., Wheeler, M. E., Schlaggar, B. L., & Petersen, S. E. (2010). Role of the anterior insula in task-level control and focal attention. Brain Structure & Function, 214(5-6), 669–680. doi:10.1007/s00429-010-0260-2

Ni, J. G., Wunderle, T., Lewis, C. M., Desimone, R., Diester, I., & Fries, P. (2016). Gamma-rhythmic gain modulation. Neuron, 92(1), 240–251. doi:10.1016/j.neuron.2016.09.003

Nolan, H., Whelan, R., & Reilly, R. B. (2010). FASTER: Fully automated statistical thresholding for EEG artifact rejection. Journal of Neuroscience Methods, 192(1), 152–162. doi:10.1016/j.jneumeth.2010.07.015

O’Connell, R. G., Dockree, P. M., & Kelly, S. P. (2012). A supramodal accumulation-to-bound signal that determines perceptual decisions in humans. Nature Neuroscience, 15(12), 1729–1735. doi:10.1038/nn.3248

O’Reilly, R. C., Wyatte, D. R., & Rohrlich, J. (2017). Deep predictive learning: A comprehensive model of three visual streams. Retrieved from doi:arXiv:1709.04654

Oldfield, R. C. (1971). The assessment and analysis of handedness: The edinburgh inventory. Neuropsychologia, 9(1), 97–113. doi:10.1016/0028-3932(71)90067-4

Oostenveld, R., Fries, P., Maris, E., & Schoffelen, J. M. (2011). Fieldtrip: Open source software for advanced analysis of MEG, EEG, and invasive electrophysiological data. Computational Intelligence and Neuroscience. doi:10.1155/2011/156869

Oostenveld, R., & Praamstra, P. (2001). The five percent electrode system for high-resolution EEG and ERP measurements. Clinical Neurophysiology, 112(4), 713–719. doi:10.1016/S1388-2457(00)00527-7

Palva, S., & Palva, J. M. (2007). New vistas for alpha-frequency band oscillations. Trends in Neurosciences, 30(4), 150–158. doi:10.1016/j.tins.2007.02.001

Parkes, L., Fulcher, B., Yucel, M., & Fornito, A. (2018). An evaluation of the efficacy, reliability, and sensitivity of motion correction strategies for resting-state functional MRI. Neuroimage, 171, 415–436. doi:10.1016/j.neuroimage.2017.12.073

Pelli, D. G. (1997). The VideoToolbox software for visual psychophysics: Transforming numbers into movies. Spatial Vision, 10(4), 437–442. doi:10.1163/156856897x00366

Pergola, G., Danet, L., Pitel, A. L., Carlesimo, G. A., Segobin, S., Pariente, J.,…Barbeau, E. J. (2018). The regulatory role of the human mediodorsal thalamus. Trends in Cognitive Sciences, 22(11), 1011–1025. doi:10.1016/j.tics.2018.08.006

Perrin, F., Pernier, J., Bertrand, O., & Echallier, J. F. (1989). Spherical splines for scalp potential and current-density mapping. Electroencephalography and Clinical Neurophysiology, 72(2), 184–187. doi:10.1016/0013-4694(89)90180-6

Peterson, E. J., & Voytek, B. (2017). Alpha oscillations control cortical gain by modulating excitatory-inhibitory background activity. bioRxiv.

Pettine, W. W., Louie, K., Murray, J. D., & Wang, X.-J. (2020). Hierarchical network model excitatory-inhibitory tone shapes alternative strategies for different degrees of uncertainty in multi-attribute decisions. bioRxiv.

Pfeffer, T., Avramiea, A. E., Nolte, G., Engel, A. K., Linkenkaer-Hansen, K., & Donner, T. H. (2018). Catecholamines alter the intrinsic variability of cortical population activity and perception. Plos Biology, 16(2), e2003453. doi:10.1371/journal.pbio.2003453

Podvalny, E., Noy, N., Harel, M., Bickel, S., Chechik, G., Schroeder, C. E.,…Malach, R. (2015). A unifying principle underlying the extracellular field potential spectral responses in the human cortex. Journal of Neurophysiology, 114(1), 505–519. doi:10.1152/jn.00943.2014

Poo, C., & Isaacson, J. S. (2009). Odor representations in olfactory cortex: "Sparse" coding, global inhibition, and oscillations. Neuron, 62(6), 850–861. doi:10.1016/j.neuron.2009.05.022

Posner, M. I., & Rothbart, M. K. (2007). Research on attention networks as a model for the integration of psychological science. Annual Review of Psychology, 58, 1–23. doi:10.1146/annurev.psych.58.110405.085516

Power, J. D., Mitra, A., Laumann, T. O., Snyder, A. Z., Schlaggar, B. L., & Petersen, S. E. (2014). Methods to detect, characterize, and remove motion artifact in resting state fMRI. Neuroimage, 84, 320–341. doi:10.1016/j.neuroimage.2013.08.048

Rafal, R. D., & Posner, M. I. (1987). Deficits in human visual spatial attention following thalamic lesions. Proc Natl Acad Sci U S A, 84(20), 7349–7353. doi:10.1073/pnas.84.20.7349

Ratcliff, R. (1978). Theory of memory retrieval. Psychological Review, 85(2), 59–108. doi:10.1037//0033-295x.85.2.59

Ratcliff, R., & McKoon, G. (2008). The diffusion decision model: Theory and data for two-choice decision tasks. Neural Computation, 20(4), 873–922. doi:10.1162/neco.2008.12-06-420

Reimer, J., Froudarakis, E., Cadwell, C. R., Yatsenko, D., Denfield, G. H., & Tolias, A. S. (2014). Pupil fluctuations track fast switching of cortical states during quiet wakefulness. Neuron, 84(2), 355–362. doi:10.1016/j.neuron.2014.09.033

Reinagel, P., Godwin, D., Sherman, S. M., & Koch, C. (1999). Encoding of visual information by lgn bursts. Journal of Neurophysiology, 81(5), 2558–2569.

Richman, J. S., & Moorman, J. R. (2000). Physiological time-series analysis using approximate entropy and sample entropy. American Journal of Physiology-Heart and Circulatory Physiology, 278(6), H2039–H2049.

Rikhye, R. V., Gilra, A., & Halassa, M. M. (2018). Thalamic regulation of switching between cortical representations enables cognitive flexibility. Nature Neuroscience, 21(12), 1753–1763. doi:10.1038/s41593-018-0269-z

Rikhye, R. V., Wimmer, R. D., & Halassa, M. M. (2018). Toward an integrative theory of thalamic function. Annual Review of Neuroscience, Vol 41, 41, 163–183. doi:10.1146/annurev-neuro-080317-062144

Roux, F., Wibral, M., Singer, W., Aru, J., & Uhlhaas, P. J. (2013). The phase of thalamic alpha activity modulates cortical gamma-band activity: Evidence from resting-state MEG recordings. Journal of Neuroscience, 33(45), 17827–17835. doi:10.1523/Jneurosci.5778-12.2013

Saalmann, Y. B., & Kastner, S. (2011). Cognitive and perceptual functions of the visual thalamus. Neuron, 71(2), 209–223. doi:10.1016/j.neuron.2011.06.027

Saalmann, Y. B., Pinsk, M. A., Wang, L., Li, X., & Kastner, S. (2012). The pulvinar regulates information transmission between cortical areas based on attention demands. Science, 337(6095), 753–756. doi:10.1126/science.1223082

Sadaghiani, S., & Kleinschmidt, A. (2016). Brain networks and alpha-Oscillations: Structural and functional foundations of cognitive control. Trends in Cognitive Sciences, 20(11), 805–817. doi:10.1016/j.tics.2016.09.004

Schiff, N. D. (2008). Central thalamic contributions to arousal regulation and neurological disorders of consciousness. Ann N Y Acad Sci, 1129, 105–118. doi:10.1196/annals.1417.029

Schmitt, L. I., Wimmer, R. D., Nakajima, M., Happ, M., Mofakham, S., & Halassa, M. M. (2017). Thalamic amplification of cortical connectivity sustains attentional control. Nature, 545(7653), 219–223. doi:10.1038/nature22073

Sherman, S. M. (2001). Tonic and burst firing: Dual modes of thalamocortical relay. Trends in Neurosciences, 24(2), 122–126. doi:10.1016/S0166-2236(00)01714-8

Shine, J. M. (2019). Neuromodulatory influences on integration and segregation in the brain. Trends in Cognitive Sciences, 23(7), 572–583. doi:10.1016/j.tics.2019.04.002

Shine, J. M., Hearne, L. J., Breakspear, M., Hwang, K., Muller, E. J., Sporns, O.,…Cocchi, L. (2019). The low-dimensional neural architecture of cognitive complexity is related to activity in medial thalamic nuclei. Neuron, 104(5), 849–855. doi:10.1016/j.neuron.2019.09.002

Siegel, M., Buschman, T. J., & Miller, E. K. (2015). Cortical information flow during flexible sensorimotor decisions. Science, 348(6241), 1352–1355. doi:10.1126/science.aab0551

Smith, A. M., Lewis, B. K., Ruttimann, U. E., Ye, F. Q., Sinnwell, T. M., Yang, Y. H.,…Frank, J. A. (1999). Investigation of low frequency drift in fMRI signal. Neuroimage, 9(5), 526–533. doi:10.1006/nimg.1999.0435

Smith, G. D., Cox, C. L., Sherman, S. M., & Rinzel, J. (2000). Fourier analysis of sinusoidally driven thalamocortical relay neurons and a minimal integrate-and-fire-or-burst model. Journal of Neurophysiology, 83(1), 588–610.

Smith, S. M., Jenkinson, M., Woolrich, M. W., Beckmann, C. F., Behrens, T. E. J., Johansen-Berg, H.,…Matthews, P. M. (2004). Advances in functional and structural MR image analysis and implementation as fsl. Neuroimage, 23, S208–S219. doi:10.1016/j.neuroimage.2004.07.051

Smulders, F. T. Y., ten Oever, S., Donkers, F. C. L., Quaedflieg, C. W. E. M., & van de Ven, V. (2018). Single-trial log transformation is optimal in frequency analysis of resting EEG alpha. European Journal of Neuroscience, 48(7), 2585–2598. doi:10.1111/ejn.13854

Song, A. H., Kucyi, A., Napadow, V., Brown, E. N., Loggia, M. L., & Akeju, O. (2017). Pharmacological modulation of noradrenergic arousal circuitry disrupts functional connectivity of the locus ceruleus in humans. Journal of Neuroscience, 37(29), 6938–6945. doi:10.1523/Jneurosci.0446-17.2017

Spaak, E., Bonnefond, M., Maier, A., Leopold, D. A., & Jensen, O. (2012). Layer-specific entrainment of gamma-band neural activity by the alpha rhythm in monkey visual cortex. Current Biology, 22(24), 2313–2318. doi:10.1016/j.cub.2012.10.020

Stitt, I., Zhou, Z. C., Radtke-Schuller, S., & Frohlich, F. (2018). Arousal dependent modulation of thalamo-cortical functional interaction. Nature Communications, 9(1), 2455. doi:10.1038/s41467-018-04785-6

Suffczynski, P., Kalitzin, S., Pfurtscheller, G., & da Silva, F. H. L. (2001). Computational model of thalamo-cortical networks: Dynamical control of alpha rhythms in relation to focal attention. International Journal of Psychophysiology, 43(1), 25–40. doi:Doi 10.1016/S0167-8760(01)00177-5

Swadlow, H. A., & Gusev, A. G. (2001). The impact of ‘bursting’ thalamic impulses at a neocortical synapse. Nature Neuroscience, 4(4), 402–408. doi:10.1038/86054

Thiele, A., & Bellgrove, M. A. (2018). Neuromodulation of attention. Neuron, 97(4), 769–785. doi:10.1016/j.neuron.2018.01.008

Tomasi, D., Chang, L., Caparelli, E. C., & Ernst, T. (2007). Different activation patterns for working memory load and visual attention load. Brain Research, 1132(1), 158–165. doi:10.1016/j.brainres.2006.11.030

Tort, A. B. L., Kramer, M. A., Thorn, C., Gibson, D. J., Kubota, Y., Graybiel, A. M., & Kopell, N. J. (2008). Dynamic cross-frequency couplings of local field potential oscillations in rat striatum and hippocampus during performance of a t-maze task. Proceedings of the National Academy of Sciences of the United States of America, 105(51), 20517–20522. doi:10.1073/pnas.0810524105

Twomey, D. M., Kelly, S. P., & O’Connell, R. G. (2016). Abstract and effector-selective decision signals exhibit qualitatively distinct dynamics before delayed perceptual reports. Journal of Neuroscience, 36(28), 7346–7352. doi:10.1523/Jneurosci.4162-15.2016

Uddin, L. Q. (2015). Salience processing and insular cortical function and dysfunction. Nature Reviews Neuroscience, 16(1), 55–61. doi:10.1038/nrn3857

Urai, A. E., Braun, A., & Donner, T. H. (2017). Pupil-linked arousal is driven by decision uncertainty and alters serial choice bias. Nature Communications, 8. doi:10.1038/ncomms14637

van Kerkoerle, T., Self, M. W., Dagnino, B., Gariel-Mathis, M. A., Poort, J., van der Togt, C., & Roelfsema, P. R. (2014). Alpha and gamma oscillations characterize feedback and feedforward processing in monkey visual cortex. Proceedings of the National Academy of Sciences of the United States of America, 111(40), 14332–14341. doi:10.1073/pnas.1402773111

van Vugt, M. K., Beulen, M. A., & Taatgen, N. A. (2019). Relation between centro-parietal positivity and diffusion model parameters in both perceptual and memory-based decision making. Brain Research, 1715, 1–12. doi:10.1016/j.brainres.2019.03.008

Vinck, M., Batista-Brito, R., Knoblich, U., & Cardin, J. A. (2015). Arousal and locomotion make distinct contributions to cortical activity patterns and visual encoding. Neuron, 86(3), 740–754. doi:10.1016/j.neuron.2015.03.028

Ward, L. M. (2013). The thalamus: Gateway to the mind. Wiley Interdisciplinary Reviews-Cognitive Science, 4(6), 609–622. doi:10.1002/wcs.1256

Waschke, L., Tune, S., & Obleser, J. (2019). Local cortical desynchronization and pupil-linked arousal differentially shape brain states for optimal sensory performance. Elife, 8. doi:10.7554/eLife.51501

Waterhouse, B. D., & Navarra, R. L. (2019). The locus coeruleus-norepinephrine system and sensory signal processing: A historical review and current perspectives. Brain Res, 1709, 1–15. doi:10.1016/j.brainres.2018.08.032

Weerda, R., Vallines, I., Thomas, J. P., Rutschmann, R. M., & Greenlee, M. W. (2006). Effects of nonspatial selective and divided visual attention on fMRI BOLD responses. Experimental Brain Research, 173(4), 555–563. doi:10.1007/s00221-006-0403-0

Weissman, D. H., Gopalakrishnan, A., Hazlett, C. J., & Woldorff, M. G. (2005). Dorsal anterior cingulate cortex resolves conflict from distracting stimuli by boosting attention toward relevant events. Cerebral Cortex, 15(2), 229–237. doi:10.1093/cercor/bhh125

Whitten, T. A., Hughes, A. M., Dickson, C. T., & Caplan, J. B. (2011). A better oscillation detection method robustly extracts EEG rhythms across brain state changes: The human alpha rhythm as a test case. Neuroimage, 54(2), 860–874. doi:10.1016/j.neuroimage.2010.08.064

Wiecki, T. V., Sofer, I., & Frank, M. J. (2013). HDDM: Hierarchical bayesian estimation of the drift-diffusion model in python. Frontiers in Neuroinformatics, 7. doi:10.3389/fninf.2013.00014

Wimmer, R. D., Schmitt, L. I., Davidson, T. J., Nakajima, M., Deisseroth, K., & Halassa, M. M. (2015). Thalamic control of sensory selection in divided attention. Nature, 526(7575), 705–709. doi:10.1038/nature15398

Wojciulik, E., & Kanwisher, N. (1999). The generality of parietal involvement in visual attention. Neuron, 23(4), 747–764. doi:10.1016/S0896-6273(01)80033-7

Wolff, M., & Vann, S. D. (2019). The cognitive thalamus as a gateway to mental representations. Journal of Neuroscience, 39(1), 3–14. doi:10.1523/Jneurosci.0479-18.2018

Wöstmann, M., Alavash, M., & Obleser, J. (2019). Alpha oscillations in the human brain implement distractor suppression independent of target selection. Journal of Neuroscience, 39(49), 9797–9805. doi:10.1523/JNEUROSCI.1954-19.2019

Wright, N. F., Vann, S. D., Aggleton, J. P., & Nelson, A. J. D. (2015). A critical role for the anterior thalamus in directing attention to task-relevant stimuli. Journal of Neuroscience, 35(14), 5480–5488. doi:10.1523/Jneurosci.4945-14.2015

Yang, G. J., Murray, J. D., Wang, X. J., Glahn, D. C., Pearlson, G. D., Repovs, G.,…Anticevic, A. (2016). Functional hierarchy underlies preferential connectivity disturbances in schizophrenia. Proceedings of the National Academy of Sciences of the United States of America, 113(2), E219–E228. doi:10.1073/pnas.1508436113

Yizhar, O., Fenno, L. E., Prigge, M., Schneider, F., Davidson, T. J., O’Shea, D. J.,…Deisseroth, K. (2011). Neocortical excitation/inhibition balance in information processing and social dysfunction. Nature, 477(7363), 171–178. doi:10.1038/nature10360

Yu, A. J., & Dayan, P. (2005). Uncertainty, neuromodulation, and attention. Neuron, 46(4), 681–692. doi:10.1016/j.neuron.2005.04.026

Zanto, T. P., Rubens, M. T., Thangavel, A., & Gazzaley, A. (2011). Causal role of the prefrontal cortex in top-down modulation of visual processing and working memory. Nature Neuroscience, 14(5), 656–U156. doi:10.1038/nn.2773

Zerbi, V., Floriou-Servou, A., Markicevic, M., Vermeiren, Y., Sturman, O., Privitera, M.,…Bohacek, J. (2019). Rapid reconfiguration of the functional connectome after chemogenetic locus coeruleus activation. Neuron, 103(4), 702–718. doi:10.1016/j.neuron.2019.05.034

## Supplementary References

Benwell, C. S. Y., Tagliabue, C. F., Veniero, D., Cecere, R., Savazzi, S., & Thut, G. (2017). Prestimulus EEG power predicts conscious awareness but not objective visual performance. Eneuro, 4(6). doi:10.1523/ENEURO.0182-17.2017

Cavanagh, J. F., Wiecki, T. V., Cohen, M. X., Figueroa, C. M., Samanta, J., Sherman, S. J., & Frank, M. J. (2011). Subthalamic nucleus stimulation reverses mediofrontal influence over decision threshold. Nature Neuroscience, 14(11), 1462–1467. doi:10.1038/nn.2925

Limbach, K., & Corballis, P. M. (2016). Prestimulus alpha power influences response criterion in a detection task. Psychophysiology, 53(8), 1154–1164. doi:10.1111/psyp.12666

Lui, K. K., Nunez, M. D., Cassidy, J. M., Vandekerckhove, J., Cramer, S. C., & Srinivasan, R. (2018). Timing of readiness potentials reflect a decision-making process in the human brain. bioRxiv.

Morgan, S. T., Hansen, J. C., & Hillyard, S. A. (1996). Selective attention to stimulus location modulates the steady-state visual evoked potential. Proceedings of the National Academy of Sciences of the United States of America, 93(10), 4770–4774. doi:10.1073/pnas.93.10.4770

Muller, M. M., Andersen, S., Trujillo, N. J., Valdes-Sosa, P., Malinowski, P., & Hillyard, S. A. (2006). Feature-selective attention enhances color signals in early visual areas of the human brain. Proceedings of the National Academy of Sciences of the United States of America, 103(38), 14250–14254. doi:10.1073/pnas.0606668103

Nunez, M. D., Vandekerckhove, J., & Srinivasan, R. (2017). How attention influences perceptual decision making: Single-trial EEG correlates of drift-diffusion model parameters. Journal of Mathematical Psychology, 76, 117–130. doi:10.1016/j.jmp.2016.03.003

O’Connell, R. G., Dockree, P. M., & Kelly, S. P. (2012). A supramodal accumulation-to-bound signal that determines perceptual decisions in humans. Nature Neuroscience, 15(12), 1729–+. doi:10.1038/nn.3248

Sheremata, S. L., Somers, D. C., & Shomstein, S. (2018). Visual short-term memory activity in parietal lobe reflects cognitive processes beyond attentional selection. Journal of Neuroscience, 38(6), 1511–1519. doi:10.1523/Jneurosci.1716-17.2017

Todd, J. J., & Marois, R. (2004). Capacity limit of visual short-term memory in human posterior parietal cortex. Nature, 428(6984), 751–754. doi:10.1038/nature02466

Zhigalov, A., Herring, J. D., Herpers, J., Bergmann, T. O., & Jensen, O. (2019). Probing cortical excitability using rapid frequency tagging. Neuroimage, 195, 59–66. doi:10.1016/j.neuroimage.2019.03.056

